# Analysis of a photosynthetic cyanobacterium rich in internal membrane systems via gradient profiling by sequencing (Grad-seq)

**DOI:** 10.1101/2020.07.02.184192

**Authors:** Matthias Riediger, Philipp Spät, Raphael Bilger, Karsten Voigt, Boris Maček, Wolfgang R. Hess

## Abstract

Regulatory sRNAs in photosynthetic cyanobacteria have been reported, but the lack of plausible RNA chaperones involved in this regulation appears enigmatic. Here, we analyzed the full ensemble of cellular RNAs and proteins using gradient profiling by sequencing (Grad-seq) in *Synechocystis* 6803. Complexes with overlapping subunits such as the CpcG1-type versus the CpcL-type phycobilisomes or the PsaK1 versus PsaK2 photosystem I pre(complexes) could be distinguished supporting a high quality of the approach. Clustering of the in-gradient distribution profiles followed by several additional criteria yielded a short list of potential RNA chaperones that include a YlxR homolog and a cyanobacterial homolog of the KhpA/B complex. The data suggest previously undetected complexes between accessory proteins and CRISPR-Cas systems, such as a Csx1-Csm6 ribonucleolytic defense complex. Moreover, the exclusive association of either RpoZ or 6S RNA with the core RNA polymerase complex and the existence of a reservoir of inactive sigma-antisigma complexes is suggested. The *Synechocystis* Grad-seq resource is available online at https://sunshine.biologie.uni-freiburg.de/GradSeqExplorer/, providing a comprehensive resource for the functional assignment of RNA-protein complexes and multisubunit protein complexes in a photosynthetic organism.

**One-sentence summary:** We provide the first global analysis of a cyanobacterium using Grad-seq, providing a comprehensive resource for the in-depth analysis of the complexome in a photosynthetic organism.

## Introduction

Non-coding (nc)RNAs constitute a major component of the transcriptional output in all classes of organisms (Cech and Steitz, 2014; Morris and Mattick, 2014). In photosynthetic cyanobacteria as well as in other bacteria, complex regulatory networks have been identified that include small regulatory ncRNAs (sRNAs) as major players in the posttranscriptional control of gene expression (Wagner and Romby, 2015; Kopf and Hess, 2015). While some sRNAs as well as cis-transcribed antisense RNAs (asRNAs) may function independent of proteins, the vast majority of ncRNAs require interactions with specific RNA binding proteins (RBPs) (Melamed et al., 2020; Quendera et al., 2020; Holmqvist and Vogel, 2018).

Cyanobacteria form a single phylum of species with very different morphologies and highly diverse lifestyles. Their genome sizes vary by approximately one order of magnitude. Cyanobacteria are monophyletic and the only bacteria that perform oxygenic photosynthesis. They are the evolutionary ancestors of chloroplasts, are of paramount ecological importance as primary producers and of high biotechnological interest (Hagemann and Hess, 2018; Vijay et al., 2019). In the unicellular model cyanobacterium *Synechocystis* sp. PCC 6803 (from here *Synechocystis*), three CRISPR systems (Scholz et al., 2013), a 6S RNA homolog (Heilmann et al., 2017) and hundreds of putative sRNAs have been identified (Kopf and Hess, 2015; Kopf et al., 2014). Some of these sRNAs are important posttranscriptional regulators. The sRNA PsrR1 caps the expression of several photosynthesis proteins and pigment biosynthesis proteins during the acclimation of the photosynthetic machinery to high light conditions (Georg et al., 2014). Likewise, the sRNA IsaR1 contributes to the adjustment of iron-sulfur cluster biosynthesis, photosynthetic electron transport and photosystem (PS) I gene expression in the acclimation response to low iron (Georg et al., 2017). Moreover, two different types of ncRNAs, the sRNA NsiR4 and the glutamine type I riboswitch, were found to control cyanobacterial nitrogen assimilation in ways that differ considerably from the archetypical model developed for *E. coli* (Klähn et al., 2015, 2018). Recently, the RBPs Rbp2 and Rbp3 (Ssr1480 and Slr0193) have been discovered to impact the thylakoid association of mRNAs encoding core subunits of both photosystems (Mahbub et al., 2020). However, despite the observed abundance of riboregulatory processes, no RBPs functionally similar to ProQ, CsrA and Hfq could be identified in cyanobacteria. While there are no candidates for the former two, a structural homolog of Hfq is present in several cyanobacteria, including *Synechocystis* (Dienst et al., 2008; Bøggild et al., 2009). However, compared to the homologs in proteobacteria, cyanobacterial Hfq is truncated, does not bind RNA and may function through protein:protein interactions (Schuergers et al., 2014).

A recently developed method uses gradient profiling by sequencing (Grad-seq) to directly detect groups of RNAs and comigrating proteins. In this approach, whole-cell lysates are fractionated on a sucrose density gradient by ultracentrifugation. The fractions are subject to mass spectrometry-based proteome measurements and RNA-sequencing analyses, hence guiding the potential discovery of new globally acting RBPs and polypeptides acting together in large multiprotein complexes (Smirnov et al., 2016). This approach has been prolific in the discovery of ProQ as a previously overlooked major small RNA-binding protein in the bacterial pathogen *Salmonella enterica* (Smirnov et al., 2016; Westermann et al., 2019) and other enteric bacteria (Melamed et al., 2020), in the discovery of FopA as a new member of the emerging family of FinO/ProQ-like RBPs (Gerovac et al., 2020), in the characterization of the seemingly noncoding RNA RyeG as an mRNA that encodes a small toxic protein (Hör et al., 2020a) and identified the unexpected involvement of exoribonuclease activity in the stabilization and activation of sRNAs in the Gram-positive pathogen *Streptococcus pneumonia* (Hör et al., 2020b).

While there are reports on the composition of protein complexes (Xu et al., 2020) and the association of these complexes with the thylakoid or cell membrane systems in the model *Synechocystis* (Baers et al., 2019), no information on RNA/protein complexes is currently available for any cyanobacterium. Here, we report the results of Grad-seq analysis applied to *Synechocystis*, identifying the sedimentation profiles of thousands of transcripts and proteins simultaneously.

We applied a hierarchical clustering approach and provide a database that supports further analyses of this dataset based on multiple filter criteria. In contrast to previous Grad-Seq studies that initially focused on the sedimentation profiles of ncRNAs followed by further experimental analysis, we directed our efforts to the co-migration analysis of proteins and RNAs. We computed a support vector machine (SVM) score for the prediction of RNA-binding proteins among the co-sedimenting proteins using RNApred (Kumar et al., 2011). In addition, we assumed that relevant RBPs would be more widely phylogenetically conserved and tested for the presence of putative homologs in a set of 57 different cyanobacteria, *Arabidopsis thaliana, E. coli* and *Salmonella enterica* and evaluated the synteny among 34 different cyanobacteria. This strategy allowed the delineation of RBPs or other proteins of interest directly. The results provide the first RNA/protein complexome resource for a photosynthetic cyanobacterium.

## Results

### Grad-seq analysis resolves the proteome and transcriptome of a photosynthetic bacterium

Triplicate cultures of *Synechocystis* were cultivated at moderate light intensities in BG11 medium. Whole cell lysates were prepared and fractionated by density gradient centrifugation as described (Smirnov et al., 2016) but using DDM as membrane solubilizer and building the gradients from sucrose. The gradients showed the colorful separation of the different pigment-containing complexes (**Figure 1A)**. After separation, the eluted fractions from each gradient were analyzed via mass spectrometry (MS)-based proteomics and RNA sequencing for the identification of their protein and RNA composition. The overall in-gradient distribution of RNA profiles (**Supplementary Figure S1A**) and protein profiles (**Supplementary Figure S1B**) showed a strong correlation (median RNA-seq r = 0.76, median MS r = 0.85) between the replicates and confirmed good reproducibility. The fractionated proteins and RNAs were also visualized using conventional denaturing polyacrylamide gels (**Figure 1A** and **B**).

**Figure 1.**
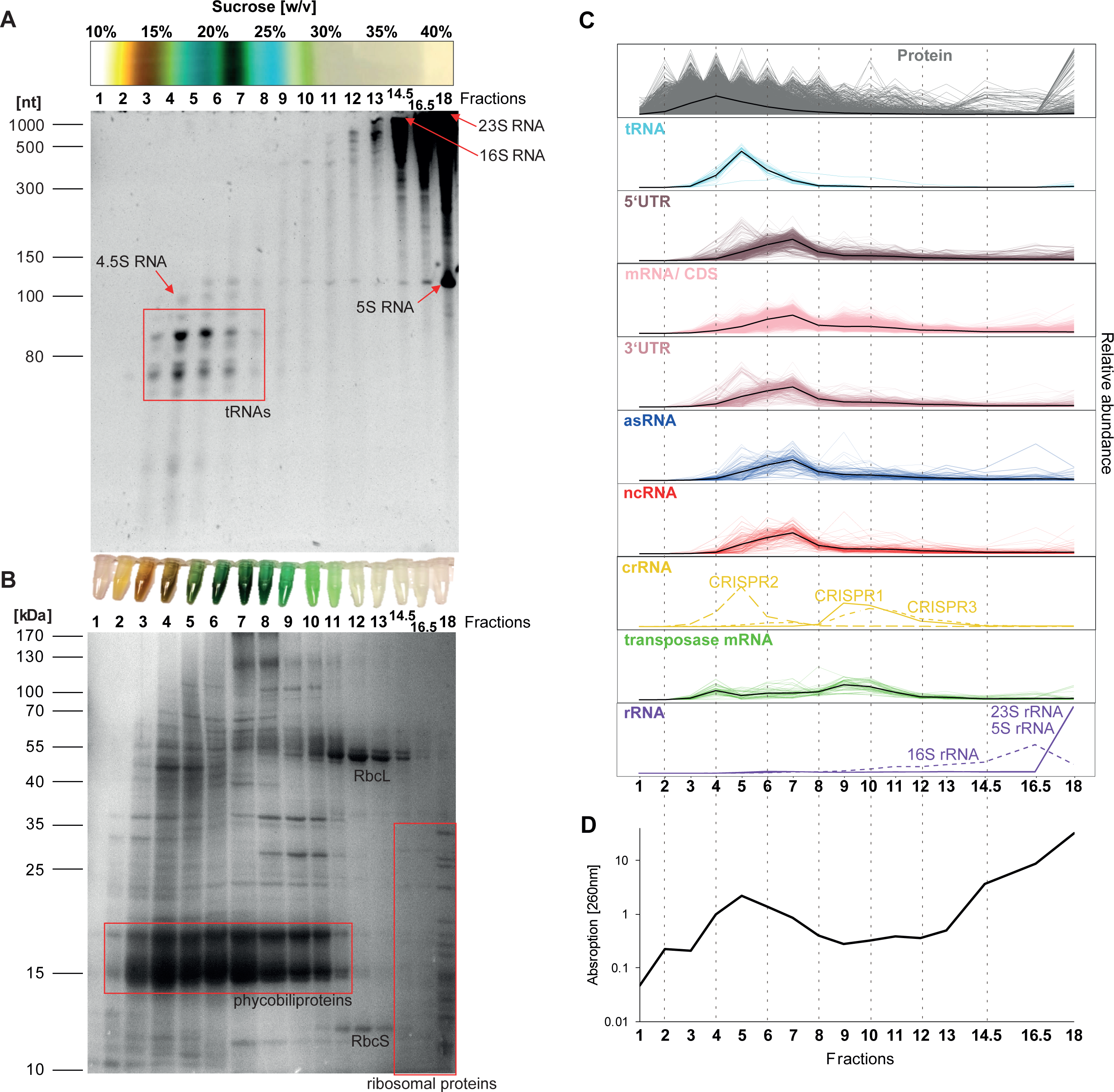
Fractionation of cyanobacterial proteins and RNAs by Grad-Seq analysis. **(A)** Top part: Gradient tube following centrifugation. The different colors match known pigment-protein complexes, containing yellow-orange carotenoids, light blue phycobilins, green chlorophyll and white material at the bottom of the tube. The different sucrose concentrations are given in % (m/v). One representative tube out of three is shown. Lower part: Separation of RNA in a 10% urea-PAA gel. The position of abundant RNA species is given for orientation. The low-range ssRNA ladder (NEB) served as a size marker (numbers to the left in nt). **(B)** Top part: Fractions after elution into collection tubes. Based on the low sample complexity of fractions 14 to 17 determined in pre-experiments, fractions 14 and 15 as well as 16 and 17 were pooled before preparing samples for mass spectrometry and RNA sequencing. Lower part: Separation of proteins in a 15% SDS-PAA gel. The positions of abundant phycobiliproteins (phycocyanin and allophycocyanin), small and large RuBisCO subunits (RbcS and RbcL) and ribosomal proteins are given for orientation. The PageRuler protein ladder (Thermo Fisher) served as a molecular mass marker (numbers to the left in kDa). **(C)** Characteristic sedimentation profiles of the entire set of proteins and of major RNA classes. **(D)** Absorption of each fraction at 260 nm measured for the purified RNA with Nanodrop. For the histograms of the Spearman correlation coefficients from the comparison of gradient profiles between the replicates see **Supplementary Figure S1**.

Following the identification of proteins and RNAs by MS and RNA sequencing, we determined the sedimentation profile of 2,394 proteins and 4,251 different transcripts detected in all three replicates (**Figure 1C)**. These transcripts consisted of 2,968 mRNAs, 530 separate 5’UTRs, 359 3’UTRs, 140 ncRNAs, 139 asRNAs, 65 transposase-associated RNAs, 42 tRNAs, 6 rRNAs and 3 types of crRNAs. A total of 3,559 annotated protein-coding genes were previously defined in this strain of *Synechocystis* (Trautmann et al., 2012); hence, we detected 67.3% of all annotated proteins. When including 122 more recently identified additional genes, we detected 65% of the 3,681 total proteins. The 4,251 different detected transcripts corresponded to 2,544 previously defined transcriptional units (TUs) out of a total of 4,091 (Kopf et al., 2014). Hence, 62.2% of all TUs were detected. In our previous definition of TUs, we assigned multiple genes and their UTRs to one and the same TU if the transcriptomic coverage did not indicate interrupting terminators or internal transcriptional start sites. However, aiming at maximum sensitivity, we kept all genes and UTRs separate here. The percentages of 67.3% detected proteins and of 62.2% detected TUs are quite exhaustive because the samples were from a single culture condition, while certain genes are expressed only under specific conditions (Kopf et al., 2014). An overview on the fraction complexity is given in **Supplementary Table S1** and the detailed distribution of all detected proteins and transcripts can be found in **Supplementary Table S2**. With our database available at https://sunshine.biologie.uni-freiburg.de/GradSeqExplorer/, we provide a comprehensive resource to scan for all detected proteins and RNAs of this dataset.

### Hierarchical cluster analysis splits the Grad-seq dataset into protein- and RNA-dominated main branches

Based on the distribution along the gradient, proteins and transcripts were assigned to 17 different clusters using the dynamic tree cut algorithm from the WGCNA R package (Langfelder and Horvath, 2008, 2012) (**Figure 2A, Supplementary Figure S2A**). A striking result from this approach was the division of the entire dataset into a protein-dominated branch whose members mainly sedimented in the lower molecular fractions and an RNA-dominated branch whose members belonged predominantly to the higher molecular fractions (**Supplementary Figure S2B** and **C**). The protein-dominated branch, represented by clusters 1 to 9, consisted of soluble proteins, small and medium protein complexes and tRNAs in lower molecular fractions 2 to 6 [∼10 - 20% sucrose]. Approximately 80% of all detected proteins were classified into this branch. The RNA-dominated branch, represented by clusters 10 to 17, included most transcripts and large protein complexes, predominantly in higher molecular fractions 6 to 18 [∼20 - 40% sucrose], and comprised ∼90% of all detected transcripts (**Figure 2B**). As expected, the sedimentation properties of different proteins varied largely and relied on the molecular weight or on the size of the associated protein complex. According to the calibration by known complexes, proteins peaking in fractions above 5 were associated with larger complexes of more than 250 kDa in size (**Supplementary Figure S3A**). In contrast, the majority of RNAs exhibited a largely similar sedimentation pattern, accumulating mainly in clusters 10, 13 and 14. Notable exceptions were RNase P RNA, tmRNA, 6S RNA as well as 5S, 16S and 23S rRNA, which sedimented towards higher buoyancies. These results are in accordance with previous observations (Smirnov et al., 2016) that RNA sedimentation profiles are largely independent of the transcript length but rely more on their corresponding binding protein[s] (**Supplementary Figure S3B**).

**Figure 2.**
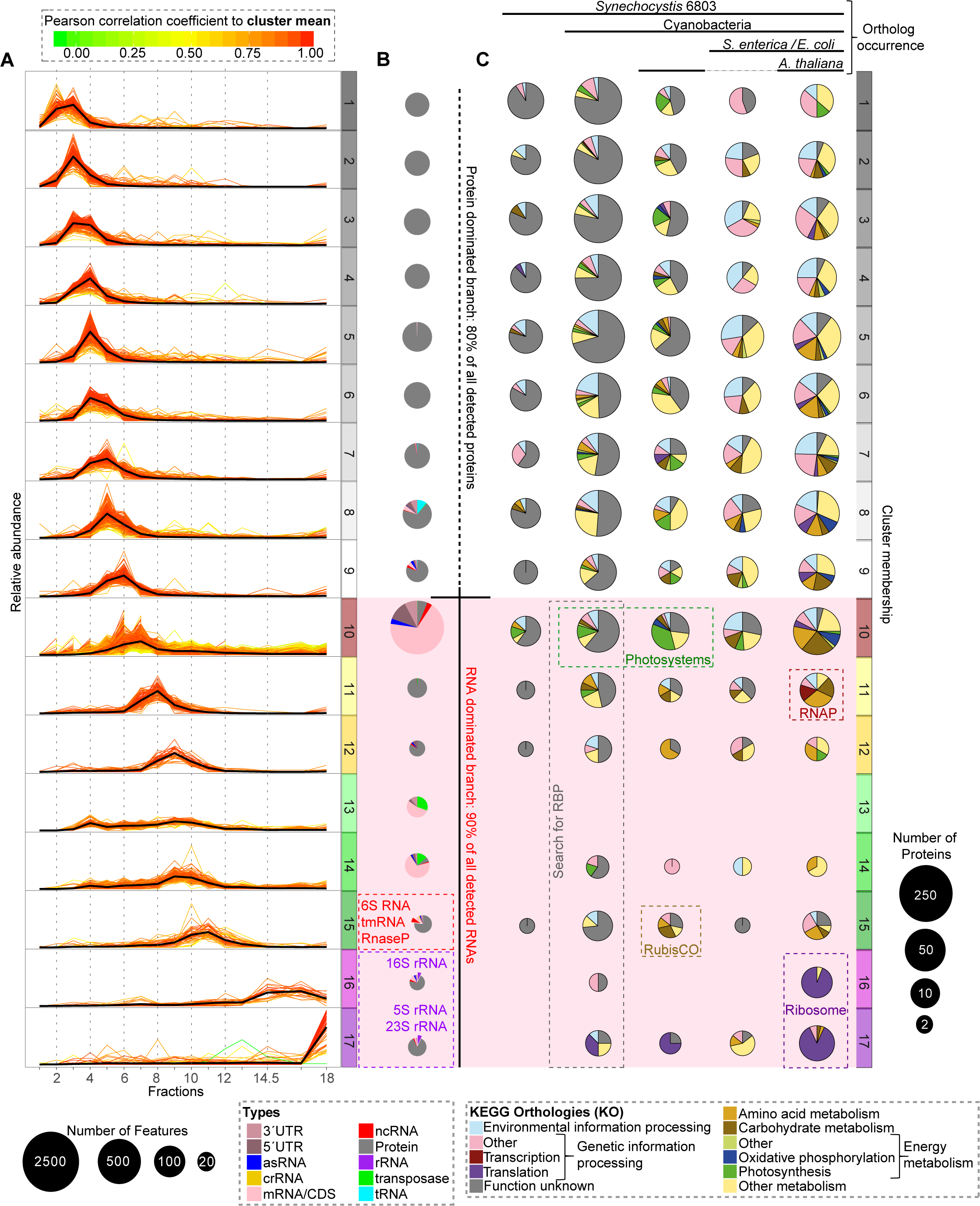
Composition of cluster profiles, functional and phylogenetic analysis. **(A)** Profiles of clusters 1 to 17 (numbers to the right) along the different fractions 1 to 18 (numbers at the X axis). The grayscaled clusters 1-9 represent the protein-dominated branch, while the colored clusters 10-17 represent the RNA-dominated branch (red background, see **Supplementary Figure S2**). The red cluster 10 represents the main RNA cluster, the yellow clusters 11-12 represent the small clusters between the RNA clusters, the green clusters 13-15 represent the other RNA clusters containing transposase-associated RNAs and housekeeping ncRNAs, such as 6S RNA, tmRNA or RNase P, while the purple clusters 16-17 represent the main ribosomal RNA clusters. **(B)** Pie charts illustrating the relative content in proteins and different RNA species of each cluster. The circle sizes correlate with the respective number of different elements in them (scale in lower left corner). **(C)** Pie charts illustrating the association of proteins in each cluster with a functional category (Kanehisa and Goto, 2000; Kanehisa, 2019; Kanehisa et al., 2019). Proteins not included in the KEGG database or annotated as “Function unknown” or “General function prediction only” were manually annotated as “Function unknown”. The phylogenetic distribution of likely orthologs was determined based on the domclust algorithm of the Microbial Genome Database (Uchiyama et al., 2019) and 59 selected genomes (**Supplementary Table S3**). As some examples, the RNAP subunits, most ribosomal and photosystem subunits or the RubisCO cluster together based on their sedimentation profiles and group together based on their phylogenetic occurrence. Largely uncharacterized proteins assigned to the RNA-dominated branch and not conserved in *Arabidopsis, Salmonella* and *E. coli* seem promising candidates as potential cyanobacterial RNA chaperones (highlighted by the dashed rectangle). The circle sizes correlate with the respective number of different proteins in them (scale to the right lower end). The distribution of homologs in the selected reference organisms is indicated by red (absence) or green (presence) rectangles at the bottom of the panel. For the visualization of sedimentation velocity versus molecular weight, see **Supplementary Figure S3**.

The proteins of our dataset were categorized into functional groups (**Figure 2C**), defined by KEGG (Kanehisa, 2019; Kanehisa et al., 2019; Kanehisa and Goto, 2000). We then compared every single protein with the presence of likely orthologs in selected other organisms (**Supplementary Table S3**). We chose a set of 56 different cyanobacterial genomes to judge the proteins conserved in cyanobacteria only, in the *E. coli* and *S. enterica* genomes combined as reference for bacteria with characterized sRNA chaperones and in the *A. thaliana* genome to detect proteins more widely conserved in the green lineage.

This led to classifying the proteins into five categories: “*Synechocystis* only”, “cyanobacteria only”, “bacteria”, “cyanobacteria and *Arabidopsis*” and “all”. The dominance of proteins of unknown function in the “*Synechocystis* only” category and their proportion of more than 50% in the “cyanobacteria only” category illustrate that these organisms are still largely unexplored. Only 20% of the *Synechocystis* proteome was assigned to clusters of the RNA-dominated branch (clusters 10 to 17 with maximum peaks at F6 to F18). These clusters included proteins that are part of large, well-known complexes, such as the majority of PS subunits, RubisCO, RNAP, most ribosomal proteins and a small number of largely undescribed proteins, depicting promising candidates for interaction with such complexes or potential RBPs, especially if not conserved in *E. coli, S. enterica* and *A. thaliana* (**Figure 2C**).

### Many transcripts show bimodal sedimentation profiles

Most transcripts were assigned to one of the three clusters 10, 13 or 14. The transcripts of clusters 13 and 14 had largely similar sedimentation profiles and were analyzed together, while the sedimentation profiles of cluster 10 transcripts clearly differed. The transcripts of cluster 10 mainly occurred in fractions 5 to 7 but with a 2^nd^ occurrence above fraction 8, often peaking around fraction 9. Therefore, the transcripts of cluster 10 were grouped according to their main peaks, yielding three subgroups (**Figure 3A**). In these subgroups, the main peaks of mRNAs, their associated asRNAs, and, as far as we know, their respective regulatory ncRNAs largely overlapped. Hence, mRNA:asRNA and mRNA:sRNA hybrids could exist in these fractions. However, the secondary peaks for mRNA occurrences did not overlap with asRNA or ncRNA peaks. Hence, the respective mRNAs cofractionated with their putative RNA regulators in the primary peak fractions and fractionated separately in the secondary peak fractions. The transcripts of clusters 13 and 14 were grouped accordingly. Transposase mRNAs are a major constituent of these clusters. Notably, only two annotated ncRNAs, Ncr0560 and Ncr1310, are included in cluster 14, exhibiting a single peak in fraction 9. Ncr0560 originates from a transcriptional start site internal to gene *slr2062* encoding an IS200/IS605 TnpB protein and therefore is closely connected to transposases. However, most transcripts of clusters 13 and 14 occurred in a narrow range of fractions 9 to 10, co-sedimenting with RNAP subunits, but also with secondary peaks below fraction 8, often in fraction 4 where they were co-sedimenting with Rps1a and Rps1b (**Figure 3B** and **Figure 4A**). Therefore, the vast majority of *Synechocystis* transcripts seem to occur in at least two states, either bound to a large complex or as unbound RNA, probably protected from degradation by one or multiple RNA chaperones.

**Figure 3.**
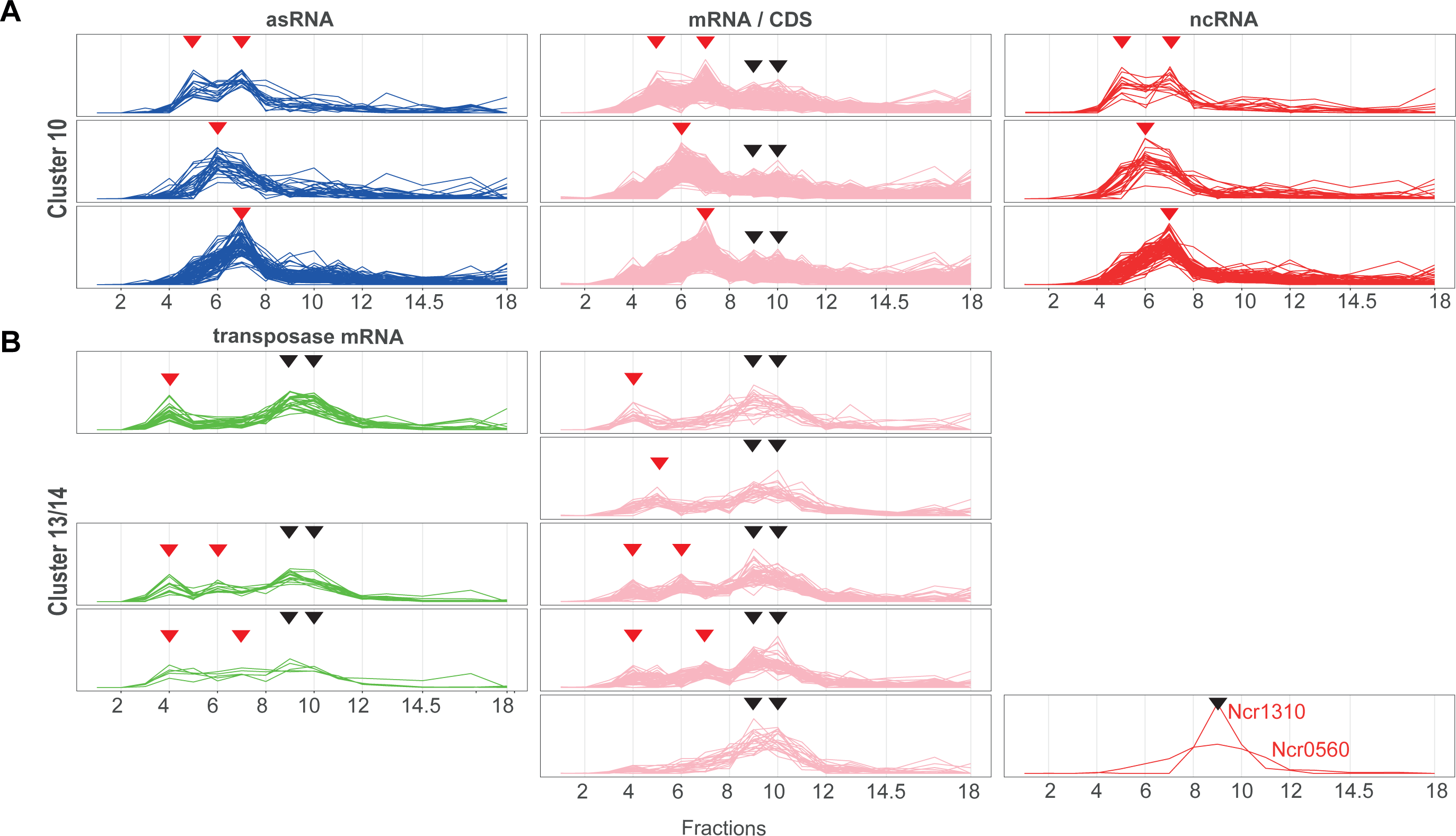
RNA sedimentation profiles. **(A)** Detailed sedimentation profiles of asRNAs (blue), mRNAs (pink) and ncRNAs (red) of cluster 10 by their peaks yield three subgroups. Transcripts with main peaks in (1) fractions 5 and 7, (2) fraction 6 or (3) fraction 7. **(B)** Detailed sedimentation profiles of transposase mRNAs (green) mRNAs (pink) and ncRNAs (red) of clusters 13 and 14 by their peaks yield five subgroups. Transcripts with secondary peaks in (1) fraction 4, (2) fraction 5, (3) fraction 4 and 6, (4) fraction 4 and 7 or (5) no secondary peaks in those fractions. The peaks relevant for the subgrouping are indicated with a red arrow, and the other peaks with a black arrow.

**Figure 4.**
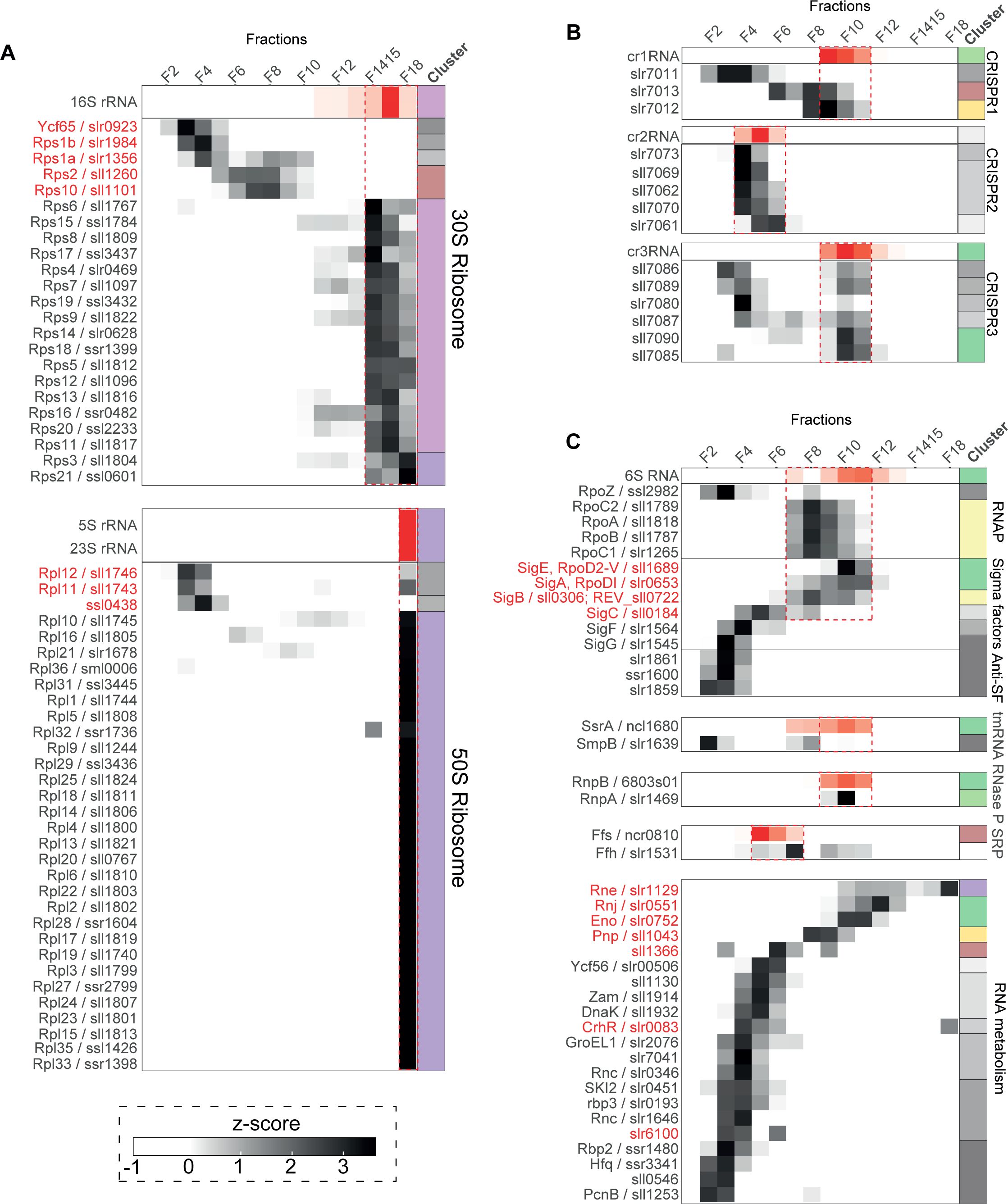
In-gradient distribution of ribonucleoprotein complexes of *Synechocystis*. The heatmap illustrates the standardized relative abundances (z-score) of each detected protein and RNA in different shades of gray (proteins) or red (RNA) alongside the fractions of the sucrose density gradient. Cluster affiliations are shown on the right (cluster color code as in **Figure 2** and **Supplementary Figure S2**). **(A)** Ribosomal RNAs and proteins. The fractions of cooccurring 16S rRNA and 30S ribosomal proteins as well as 5S, 23S rRNA and 50S ribosomal proteins are edged by a red box. The names of ribosome-associated proteins with diverging gradient profiles are indicated in red, such as Ycf65, Rps1b, Rps1A, Rps2, Rps10 (30S subunit), Rpl12, Rpl11 and Ssl0438 (50S subunit). **(B)** CRISPR RNAs and associated proteins. The fractions of respective cRNAs and cooccurring CRISPR proteins are edged by red boxes. **(C)** RNAP subunits, sigma factors and 6S RNA, essential housekeeping RNAs and their protein binding partners, as well as selected proteins involved in RNA metabolism. The fractions of cooccurring RNAP subunits, sigma factors and 6S RNA, as well as the tmRNA, RNase P RNA and signal recognition particle (SRP) RNA cooccurring with their respective cognate binding proteins, are edged by red boxes. Sigma factors and proteins involved in RNA metabolism with peaks above fraction 5 are marked in red.

### Grad-seq analysis allows the detection of ribonucleoprotein complexes

Fifty-three ribosomal or ribosome-associated proteins and the ribosomal RNAs (rRNAs) were present in the fractions containing the complexes with the highest sedimentation coefficients. A preference for the entire set of ribosomal components was restricted to fraction 18 (**Figure 4A**). Notable exceptions were Rpl11/Sll1743, Rpl12/sll1746, Rps2/sll1260, Rps10/sll1101 and Rpl16/sll1805, pointing to their association with other complexes or transcripts. Moreover, Ycf65/slr0923, previously reported as a plastid-specific ribosomal protein in some eukaryotic algae (Turmel et al., 2009) and higher plants (Yamaguchi and Subramanian, 2003), was not found to be associated with other r-proteins, nor were the Rps1a/slr1356 and Rps1b/slr1984 proteins (**Figure 4A**). Rps1a and Rps1b rather may play a role in the Shine-Dalgarno-independent initiation of translation in cyanobacteria (Nakagawa et al., 2010; Mutsuda and Sugiura, 2006). Their occurrence in fractions 3 to 5 (Rps1a) 3 to 10 (Rps1b) indicates a wide set of associations to different transcripts while the missing association to the ribosomal subunits likely was due the used buffer conditions.

CRISPR-Cas systems are particularly interesting macromolecular complexes that are known to consist of both protein and RNA components. CRISPR-Cas systems confer a form of acquired immunity against invading nucleic acids in many different bacteria and archaea and can be classified into greatly varying subtypes that belong to two major classes, with multi-subunit effector complexes in class 1 and single-protein effectors in class 2 (Makarova et al., 2020). Despite the high level of divergence among different systems, CRISPR-Cas systems usually consist of repeat-spacer arrays and the associated protein-coding *cas* genes.

*Synechocystis* encodes three separate and complete CRISPR-Cas systems that are highly expressed and active in interference assays (Scholz et al., 2013; Reimann et al., 2017; Kieper et al., 2018; Behler et al., 2018; Scholz et al., 2019). According to the presence of signature genes, these three systems were classified as subtype I-D, subtype III-D and subtype III-Bv systems (Scholz et al., 2013; Behler et al., 2018).

The gradient profiles of the crRNAs of the three CRISPR systems of *Synechocystis* varied vastly from one another. While CRISPR1 RNAs (cr1RNAs) were mainly detected in fraction 9 (cluster 14), CRISPR3 RNAs (cr3RNAs) peaked in fraction 10 (**Figure 4B**). However, the detected CRISPR2-associated proteins colocalized together with the cr2RNAs in low molecular fractions 4 to 6 at the same sedimentation level as tRNAs. The detected CRISPR1-associated proteins were scattered throughout the gradient, with only the Cas7/Csc2 protein Slr7012 cofractionating with cr1RNAs in higher molecular fractions 9 to 11. This tight association with cr1RNAs is consistent with the known function of this protein in binding the variable crRNA spacer sequence (Hrle et al., 2014). All structural proteins associated with the CRISPR3 system (proteins Cmr1, Cmr5, Cmr4, Cmr3 and Cmr2/Cas10 encoded by the genes *sll7085* to *sll7087, sll7089* and *sll7090*) fractionated together and with the CRISPR3-associated crRNAs, with a peak in fraction 10. This co-fractionation indicates that the interference complex remained largely intact during the preparation. However, those proteins exhibited a second peak in the lower fractions F3-F4 without associated cr3RNA or the Cas10 protein Sll7090, hence constituting a possibly nonfunctional subcomplex or resulting from partial disassembly of the complex during lysis preparation or centrifugation (**Figure 4B**). Interestingly, Slr7080, a Csx3 accessory protein, cofractionated with these two CRISPR3 complexes. Cas accessory genes encode diverse groups of proteins that cooccur only sporadically with CRISPR-Cas systems and are not essential for their functionality (Shmakov et al., 2018; Shah et al., 2019). Slr7080 has been characterized as a Csx3-AAA protein (Shah et al., 2019), a distant member of the Cas Rossmann fold (CARF) protein family (Makarova et al., 2014). CARF proteins sense ligands such as cyclic oligoadenylate (cOA) signaling molecules that can be synthesized by the polymerase domain of Cas10 upon recognition of its target (Kazlauskiene et al., 2017; Niewoehner et al., 2017; Han et al., 2018). The *Archaeoglobus fulgidus* Csx3 protein AfCsx3 is a ring nuclease that degrades the CRISPR signaling molecule cOA4 (Athukoralage et al., 2020). However, Slr7080/Csx3 is substantially larger than AfCsx3 (310 compared to 110 amino acids) and possesses an additional AAA ATPase domain in the C-terminal region (Shah et al., 2019). Our data suggest that Slr7080/Csx3 is expressed relatively abundantly and likely is physically linked to both the cr3RNA-free subcomplex and the complete functional interference complex.

Compelling evidence for an even more intriguing complex of Cas accessory proteins was found for the CRISPR2 system in which the four proteins Slr7061/Csx1, Sll7062/Csm6, Sll7070 and Slr7073 overlapped with the fractionation of cr2RNAs (**Figure 4B**). Slr7061/Csx1 was bioinformatically classified as a CARF-HEPN domain protein and Sll7062/Csm6 as a CARF-RelE domain-containing protein (Shah et al., 2019). The protein most closely structurally related to Slr7061/Csx1 is the *Sulfolobus islandicus* Csx1 protein, an RNase that becomes allosterically activated upon cOA4 binding (Molina et al., 2019). Csm6 is a cOA-dependent CRISPR ribonuclease as well (Garcia-Doval et al., 2020); hence, our data suggest the presence of a ribonucleolytic defense complex associated with cr2RNAs. The other two proteins, Sll7070 and Slr7073, are entirely uncharacterized, but the localization of their genes that frame the CRISPR2 *cas1cas2* adaptation module suggests a functionality closely associated with the CRISPR2 system. Prior to this work, neither the expression of Slr7061/Csx1, Sll7062/Csm6, Sll7070 and Slr7073 at the protein level nor their likely physical association with cr2RNAs was known. Hence, these proteins are suggested as highly interesting candidates for future experimental analyses.

The cyanobacterial RNAP is unique due to splitting of the β’ subunit into an N-terminal γ (RpoC1) and a C-terminal β’ (RpoC2) subunit (Schneider and Hasekorn, 1988) and a >600 amino acid insertion into RpoC2 (Iyer et al., 2004). Moreover, the interplay between the core subunits, multiple sigma factors (9 in *Synechocystis*) and inhibitory 6S RNA is only partially understood (Heilmann et al., 2017). We observed that all RNAP core subunits colocalized with the sigma factors SigB and SigC, as well as 6S RNA, SigE and SigA in higher molecular fractions, indicating the parallel presence of multiple forms of the RNAP holoenzyme. In contrast, the bulk of the RNAP ω subunit (RpoZ), as well as SigF and SigG, together with three detected anti-sigma factors was distributed only into the low molecular fractions, suggesting the presence of a reservoir of inactive binary complexes (**Figure 4C**). Furthermore, we noticed the mutually exclusive occurrence of either RpoZ or the 6S RNA together with the RNAP core subunits (see F8 in **Figure 4C**). This observation may explain the observation that cyanobacterial RpoZ facilitates the association of the primary sigma factor SigA with the RNAP core stimulating transcription of highly expressed genes (Gunnelius et al., 2014). In *E. coli*, 6S binds the β/β’ subunits (Wassarman and Storz, 2000). Hence, RpoZ may affect this binding by interacting with the γ and β’ subunits in *Synechocystis* (Gunnelius et al., 2014).

RNase P and the tmRNA (SsrA) showed similar sedimentation patterns as the RNAP holoenzyme and 6S RNA, while the signal recognition particle mostly clustered together with the bulk of mRNAs (**Figure 4C**). Several other factors, which were annotated to different functionalities in RNA metabolism, were found associated with larger complexes. Such factors were the DEAD-box RNA helicase CrhR, Sll1366 (a putative SNF2 helicase), polynucleotide phosphorylase (PNPase), enolase, RNase J and RNase E. The latter enzyme is central for RNA metabolism in bacteria because it is involved in numerous functions, such as the posttranscriptional regulation of gene expression, maturation of transcripts or initiation of mRNA degradation (Bandyra and Luisi, 2018). Examples in *Synechocystis* include the posttranscriptional regulation of gene expression with the sRNA PsrR1 as specificity factor (Georg et al., 2014), operon discoordination of the *rimO-crhR* dicistron (Rosana et al., 2020) or the maturation of functionally active crRNAs from the CRISPR3 system (Behler et al., 2018). Indeed, RNase E was found in several fractions, peaking together with ribosomal proteins, cooccurring with the bulk of mRNAs as well as cr3RNAs (**Figure 4C**). Cyanobacterial RNase E forms a complex with PNPase through an oligoarginine repeat (Zhang et al., 2014b), consistent with their co-occurrence in fraction 10 while their separate occurrences in other fractions (**Figure 4C**) indicates that the PNPase-RNase E complex is likely only one of several different combinations in which RNase E exists in the cell.

These results illustrate the sedimentation profiles of native ribonucleoprotein complexes in a sucrose density gradient, supplying valuable information about the compositional structure of such complexes in a cyanobacterial model organism. In contrast to the Grad-seq analysis performed with *Salmonella* (Smirnov et al., 2016), Hfq was in the density gradients not even close to any RNAs or known ribonucleoprotein complexes (**Figure 4C**), confirming the different functionality of this protein in cyanobacteria.

### Sedimentation profiles of mature and intermediate protein complexes associated with the thylakoid membrane provide insights into the complexity of the photosynthetic lifestyle

Focusing on protein complexes of the cyanobacterial thylakoid membrane helps to understand the factors that maintain its photosynthetic machinery. Most core components of these complexes co-sedimented, indicating the presence of largely intact PSI, PSII and cytochrome-b_6_f core complexes (**Figure 5A**), NADH-dehydrogenase (NDH-1) and cytochrome-c-oxidase complexes as well as F0F1ATP synthase (**Supplementary Figure S4A-C**). The detection of several small proteins, such as NdhP/sml0013 (Schwarz et al., 2013; Zhang et al., 2014a), NdhN, NdhH, NdhJ, and NdhO (He and Mi, 2016), co-sedimenting with other NDH-1 proteins, supports the validity of the data. The soluble electron carriers plastocyanin (PetE), cytochrome c_6_ (PetJ) and ferredoxins (PetF, Fdx) were mainly located between the lowest molecular fractions 2 and 3 and were not associated with any complex except for Fdx7 (**Supplementary Figure S4D**).

**Figure 5.**
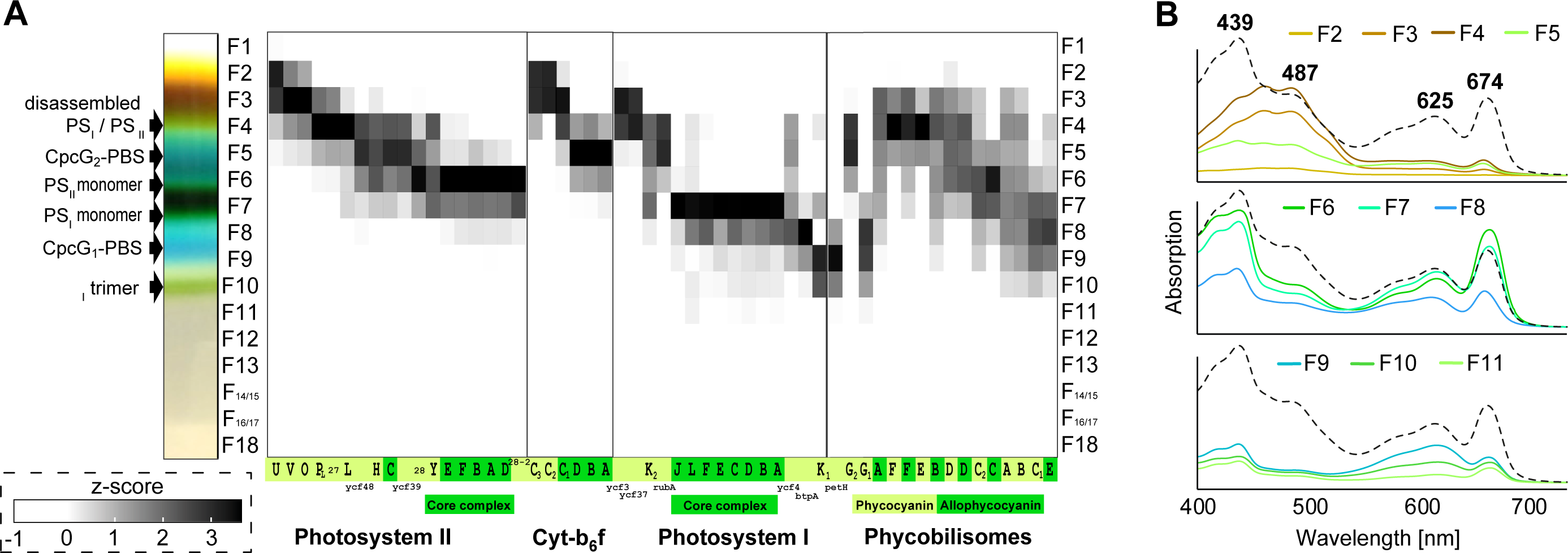
Heatmap showing the in-gradient distribution of the subunits of the major protein complexes involved in photosynthesis. **(A)** The sucrose density gradient [w / v 10 % - 40 %] showed a characteristic color pattern (left) corresponding to the pigments bound to distinct macromolecular complexes detected in the same sedimentation range (grayscale heatmap). The in gradient distribution of detected photosystem subunits and phycobiliproteins is given in the heatmap with standardized relative abundances (z-score). The profiles of CpcG1 and CpcG2 are shown separately to illustrate the difference between the G-type PBS (CpcG1 as rod-core linker) and L-type PBS (CpcG2 as rod-core linker) types (Kondo et al., 2005; Liu et al., 2019). PsbP_L_ refers to the protein Sll1418 that has been characterized as a PsbP-like protein associated with PSII (Ishikawa et al., 2005; Sveshnikov et al., 2007; Summerfield et al., 2005) **(B)** Absorption spectra taken from the indicated pigmented fractions. The dashed lines indicate the average absorption spectrum of all pigmented fractions. Complex subunits are abbreviated with the last letter of the gene name. Subunits of PSII, e.g., psb**A**, of the cytochrome-b_6_f complex, e.g., pet**A**, subunits of PSI, e.g., psa**A**, subunits of phycobilisomes, e.g., phycocyanins cpc**A** or allophycocyanins apc**A**. Green coloring indicates membership to the core complex, while green-yellow coloring refers to subcomplexes, accessorial parts of the complex or assembly factors, showing gradient profiles differing from the corresponding core complex.For the in-gradient distribution of other membrane associated complexes and proteins associated to photosynthesis see **Supplementary Figure S4**.

The occurrence of the PSII and PSI core complexes is in accordance with the measured absorption spectra for fully assembled complexes in fractions 6 and 7 (**Figure 5B**). The three membrane-extrinsic PSII subunits PsbO, PsbV and PsbU, located at the lumenal side of the thylakoid membrane, were disconnected and co-sedimented in fraction 3 (**Figure 5A**). Several other PSII accessory proteins were identified in lower fractions as well, including Psb27 (**Figure 5A**) and most PSII assembly factors (reviewed by (Nickelsen and Rengstl, 2013)) (**Supplementary Figure S4E**). A PSII subcomplex lacking PSII core reaction center polypeptides but containing Psb27 was described as a PSII repair cycle chlorophyll-protein complex (Weisz et al., 2019); therefore, corresponding complexes in transition between disassembled and assembled PSII were likely present in fractions 4 to 5. This hypothesis is supported by the measured absorption spectra, which mainly showed the presence of carotenoids but no chlorophylls (**Figure 5B**). Detected PSII assembly factors co-sedimenting with and potentially associated with PSII subcomplexes include Slr1471, PratA/Slr2048, CtpA/Slr0008 and Ycf48/Slr2034, which are involved in D1 integration, processing and stabilization in the early stage of reaction center assembly; Psb27/Slr1645 and Sll0606, which are involved in CP43 assembly and integration in the later stage of reaction center assembly and PSII repair cycle; and Slr1768, Sll1390 and Sll1414, which are involved in D1 turnover as part of the PSII repair cycle (Nickelsen and Rengstl, 2013). Interestingly, few PSII assembly factors showed a more complex sedimentation pattern with additional peaks in higher molecular fractions. Slr1768 exhibited a second peak in F11 and Slr2013 in F18, while Sll0933, the homolog of the PSII assembly factor PAM68 in *Arabidopsis* (Rast et al., 2016; Rengstl et al., 2013), solely sedimented in higher molecular fractions F11, F14 to F18, leaving their exact role in PSII assembly in *Synechocystis* open for discussion.

Furthermore, our data illustrate the roles of PsaL and PsaK in PSI trimer formation. PsaL and PsaK are the last subunits to be incorporated into PSI in the trimer assembly process, with the incorporation of PsaK2 mainly involved in state transition and PsaK1 in the constitutive state (Dühring et al., 2007; Fujimori et al., 2005), while PsaL is required for PSI trimer formation (Chitnis and Chitnis, 1993). The intact PSI monomer could be easily detected, but the PSI trimer peak was only weakly detectable, since the majority of PSI seemed to occur monomerically, possibly caused by the preparation. However, the peak appeared more clearly when the proteomics data were normalized to constant protein amounts per fraction (**Supplementary Figure S4G** and **H**). Matching previous findings (Dühring et al., 2007; Fujimori et al., 2005), our data show the co-sedimentation of PsaL and PsaK1 with the fully assembled PSI monomer as well as the trimer, while the majority of PsaK2 was only partially incorporated into the PSI monomer together with most assembly factors. Similar to previous observations of multiple peaks of some PSII assembly factors, Ycf4/Sll0226 exhibited a second and VIPP1/Sll0617 exhibited two more peaks in higher molecular fractions (**Supplementary Figure S4F**).

The sedimentation profiles of the phycobiliproteins pointed to another aspect of the complex dynamics of the photosynthetic apparatus. Two types of phycobilisomes (PBS) were previously described. The main phycobilisome (CpcG1-PBS type) is characterized by CpcG1 as a rod-core linker and the smaller CpcL-PBS type is characterized by CpcG2 as a rod core linker (Kondo et al., 2005; Liu et al., 2019). The distinction between those two types according to their protein composition illustrates the different sedimentation profiles of CpcG2-PBS and CpcG1-PBS within the pigmented sedimentation range (**Figure 5**). The association of CpcG2 with PsaK2 was suggested under high light conditions to function as a PSI antenna involved in state transitions (Kondo et al., 2007, 2009). As part of an NDH1L–PSI–CpcG2-PBS supercomplex, CpcG2 is involved in facilitating efficient cyclic electron transport (Gao et al., 2016). Interestingly, ferredoxin-NADP^+^ reductase (FNR, PetH) and PsaK1 colocalize with CpcG1-PBS. The association of FNR with phycobilisomes has previously been shown (Kondo et al., 2005; Liu et al., 2019), while the interaction of PsaK1 and CpcG1-PBS, analogous to the interaction of PsaK2 and CpcG2-PBS, has not yet been shown.

### Smaller complexes and protein-protein interactions in toxin-antitoxin systems and regulation

In addition to the analysis of ribonucleoprotein complexes and membrane-bound protein complexes, the dataset also provides interesting clues about smaller complexes consisting of just some interacting proteins.

One class of such complexes with frequently just two proteins are the toxin–antitoxin (TA) systems. TA systems consist of a stable toxic protein and its unstable cognate antitoxin. In the case of Type II TA systems, both of these components are small proteins (Schuster and Bertram, 2013). Previously, 69 Type II TA systems were predicted for *Synechocystis* (Kopfmann et al., 2016). While the majority of these proteins are plasmid-localized, consistent with their function in plasmid maintenance, 22 predicted TA systems are encoded in the main chromosome. These must have functions distinct from postsegregational killing but are severely understudied. We detected at least four such TA systems: Slr0770/Slr0771 and Ssl0258/Ssl0259 in cluster 2; Ssl2138/Sll1092 in cluster 5 and Ssl2245/Sll1130 in cluster 8 (**Figure 6A**). From these, the first two have not been studied so far, while the functionality of Ssl2138/Sll1092 as a TA system was experimentally verified (Ning et al., 2013) and for Ssl2245/Sll1130, some evidence exists that it is involved in a wider regulatory context (Srikumar et al., 2017). Interestingly, Ssl2245/Sll1130 were assigned to cluster 8, suggesting that they were part of a larger complex. Hence, our data not only support the existence of these TA systems but also show their active expression and support their supposed mode of action due to their co-fractionation.

**Figure 6.**
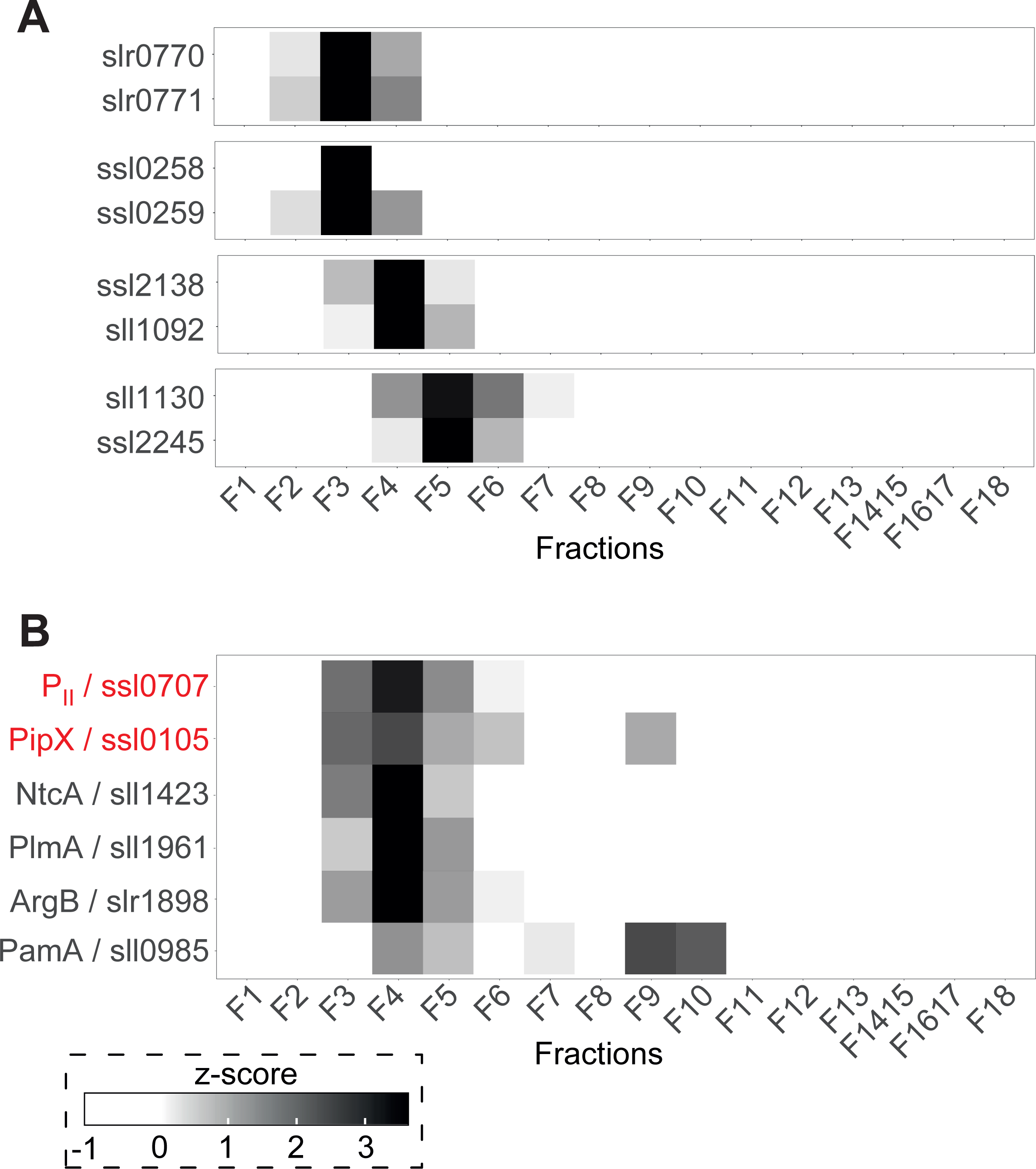
In-gradient distribution of protein-protein complexes consisting of few interacting proteins. The heatmap illustrates the standardized relative abundances (z-score), giving the in gradient distribution of **(A)** all detected TA systems and **(B)** proteins interacting with the nitrogen regulatory proteins P_II_ and PipX.

Other smaller protein complexes have regulatory functions. Here, we chose proteins interacting with components of the nitrogen regulatory system for illustration. The GntR-like transcriptional regulator PlmA is in an inactive state when bound to P_II_-PipX (Labella et al., 2016) but is involved in phycobilisome degradation when activated under nitrogen starvation (Sato et al., 2008). The formation of the P_II_-PipX complex also controls the activity of the transcriptional master regulator for nitrogen starvation, NtcA (for review see (Labella et al., 2020a)). During high nitrogen abundance, P_II_ forms a complex with either N-acetylglutamate kinase (NAGK, ArgB), which stabilizes it and therefore enhances arginine synthesis. Furthermore, it binds approximately 85% of PipX to sequester it from interacting with NtcA and thus inhibits its regulatory properties. The other 15% of PipX is bound by PlmA to keep it inactive. During low nitrogen, P_II_ is sequestered by 2OG, and NtcA-dependent gene expression is activated upon binding to PipX (Llacer et al., 2010; Forcada-Nadal et al., 2018). Our data illustrate the co-occurrence of the P_II_-PipX complex and all mentioned proteins interacting with its single components (**Figure 6B**). Another protein interacting with P_II_ during high nitrogen conditions is the membrane protein PamA, implicated in the control of genes of nitrogen and sugar metabolism (Osanai et al., 2005). Notably, this is the only one of the previously mentioned proteins that exhibited peaks outside the range of the P_II_-PipX complex, thus potentially belonging to a larger complex (**Figure 6B**). Interestingly, the single occurrence of PipX in fraction 9 (**Figure 6B**) might be linked to a recently suggested new functionality as an interactor with the translation machinery (Cantos et al., 2019).

### Classification of sedimentation zones linked to various steps of protein biosynthesis assisting in the identification of cyanobacterial RNA chaperones

Most transcripts accumulated in a narrow gradient range, equivalent to large protein complexes of ∼300 kDa in size, similar to the PS monomers. Intriguingly, this section was largely free of known RNA interacting complexes, indicating the presence of one or more RNA chaperones responsible for the characteristic sedimentation pattern. We classified these transcripts of cluster 10 as group 1 RNAs. Other transcripts assigned to clusters 13 and 14 were classified as group 2 RNAs **(Figure 7**).

**Figure 7.**
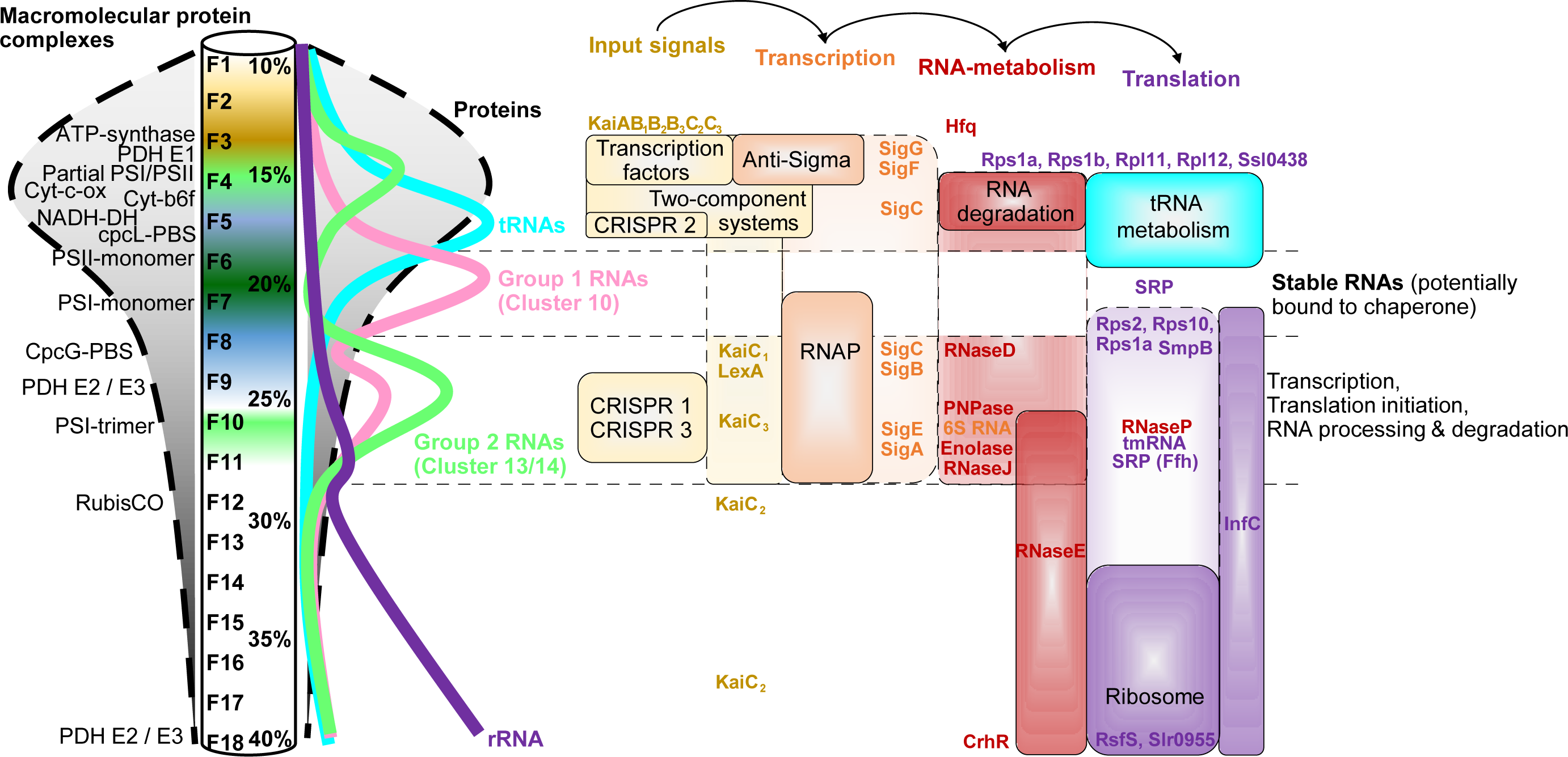
Sedimentation zones functionally linked to gene expression. The scheme illustrates the sedimentation profiles of intact macromolecular protein complexes (left) alongside the sucrose density gradient, matching the observed pigment pattern. The overall protein and RNA distribution along the gradient is depicted on the right of the gradient scheme. The majority of proteins (∼80 %) occur in the lower molecular fractions (below F6, clusters 1-9), indicating their association with small complexes or occurrence as soluble proteins. Proteins that fall into that group and are involved in gene expression are the majority of transcription factors and two component systems, anti-sigma factors, the sigma factors SigG and SigF, several proteins involved in RNA degradation and tRNA metabolism. The minority of proteins (∼20 %) occur in the higher molecular fractions (above F5, cluster 10-17), indicating their association with large complexes. Proteins that fall into that group and are involved in protein biosynthesis are the RNA polymerase (RNAP), the sigma factors SigC, SigB, SigE and SigA, several proteins and RNA-protein complexes involved in RNA metabolism, such as RNase D, RNase J, RNase E, RNase P, PNPase, Enolase, CrhR, 6S RNA, tmRNA, the signal recognition particle (SRP) and the ribosomes. A large portion of these proteins and RNA-protein complexes cooccur in fractions 8-11. Most detectable RNAs occur in fractions 8-11 as well, indicating their likely direct association with complexes composed of proteins involved in transcription, RNA degradation, RNA processing or translation initiation. Notably, most detected RNAs fall into two groups. Group one comprises those that sediment in high abundance in fractions 6-7 (cluster 10), potentially forming stable interactions with RNA chaperones. Group two includes those RNAs that generally sediment in lower abundances in fraction 4 (cluster 13-14), potentially not bound to any RNA chaperone and thus are not as stable when not directly associated in a larger complex. tRNAs are the exception, mainly sedimenting in high abundance in fraction 5 (cluster 8) together with the majority of tRNA synthetases and likely not bound to a larger complex, while rRNAs seem to be solely attached to the ribosomes.

Most RNA-interacting complexes involved in protein biosynthesis sedimented deeper, accumulating in higher molecular fractions. These proteins included the RNAP holoenzyme, 6S RNA, RNase P, tmRNA (SsrA), several other proteins functionally linked to RNA degradation and a subset of ribosomal proteins. SmpB, the cognate tmRNA RBP occurred in F2 indicating a substantial amount of free protein. However, the second peak in F7 and F8 overlapped with tmRNA indicating complex formation (**Figure 4C**). The co-occurrence of tmRNA and Rps1a in F8 and F10 (**Figure 4A**) is consistent with reports on the direct physical interaction between Rps1 and tmRNA in *E. coli* (Wower et al., 2000).

A large set of transcripts, including 16S rRNA, peaked in the same range, indicating the location of a large transcription-translation complex still binding RNAs (**Figure 7**). Proteins that sedimented in these fractions are another group of interesting candidates with potential involvement in RNA metabolism. Interestingly, although CRISPR1, CRISPR3 and most transposase mRNAs mainly co-sedimented with those complexes, their involvement in specific regulatory aspects is not known.

Based on these findings, further analyses were performed to characterize the proteins co-sedimenting with transcripts and to determine interesting candidates for future research (**Supplementary Figure S5**). We primarily focused on 19 % (451 / 2394) of all detected proteins, which were assigned to the RNA-dominated clusters in the higher molecular fractions (cluster ≥10), to which the essential complexes of protein biosynthesis also belonged. Specifically, 27 % (124 / 451) of those proteins that were annotated as unknown function were of high interest. Of those, a total of .41% (51 / 124) were conserved in at least 50 % of selected cyanobacterial genomes but not in *E. coli, S. enterica* or *A. thaliana* (**Supplementary Table S3**). Hence, these 51 proteins are conserved, largely undescribed and exhibit sedimentation profiles linking them to RNA metabolism. To aid the selection further, an support vector machine (SVM) score was computed for the prediction of RNA-binding proteins from their amino acid sequence using the RNApred web server (Kumar et al., 2011). We performed this prediction for all proteins and included it in the database associated with this paper. Approximately 27 % of the resulting 51 proteins fulfilled the SVM score criteria, which were set according to the RNApred performance for *Synechocystis* proteins (**Supplementary Figure S6**), narrowing the list of RNA chaperone candidates to only 14 proteins (0.67%) of the initially 2,394 detected proteins (**Table 1**).

**Table 1:** Shortlist of RNA chaperone candidates based on multiple filter criteria (shown in Figure 7). The locus tag is given, followed by cluster no. to which the protein was assigned, the fractions in which the abundance peaked (Max) and the fractions in which peaks occurred. The following columns give the predicted number of amino acids (AA), the relative conservation and SVM scores, and finally comments and annotation. All listed proteins were assigned to the phylogenetic group “Cyanobacteria”. For details of the taxonomic distribution and SVM scores, see **Supplementary Figure S5** and **S6**. For a synteny analysis of selected RNA chaperone candidates, see **Supplementary Figure S7**.

| Locus tag | Cluster | Max | Fractions | AA | Relative conservation | SVM score | Comments and annotation |
| --- | --- | --- | --- | --- | --- | --- | --- |
| sll1424 | 10 | 6 | 6, 10 | 491 | 1 | 0.78 | Putative SMC (structural maintenance of chromosomes) domain |
| sll1515/gifB | 10 | 6 | 6 | 149 | 0.71 | 1.97 | glutamine synthetase inactivating factor IF17 |
| slr0169 | 10 | 7 | 7 | 213 | 0.94 | 0.99 | CsoD1 ortholog (part of β-Carboxysome |
| slr0211 | 10 | 7 | 4, 7 | 403 | 0.99 | 0.83 | acyl-CoA N-acyltransferase domain |
| slr1081 | 10 | 7 | 4, 7 | 210 | 0.75 | 0.56 | DUF820 featuring a restriction endonuclease domain |
| ssl2064 | 10 | 7 | 4, 7 | 75 | 0.64 | 2.22 | DUF4160 |
| ssl2874 | 10 | 6 | 4, 6, 11 | 89 | 0.9 | 2.42 | RemA homolog; KEGG K09777 |
| sll0319 | 11 | 8 | 3, 8 | 297 | 0.57 | 1.55 | DUF3747, periplasmic protein |
| slr0287 | 12 | 9 | 5, 9 | 118 | 0.9 | 1.02 | KH domain-containing protein; KEGG K06960 |
| sll1939 | 14 | 9 | 3, 9, 11, 13 | 214 | 0.89 | 0.53 | FtsZ interacting Ftn6 homolog |
| sll0284 | 15 | 11 | 11 | 579 | 0.93 | 0.64 | DUF87 containing an N-terminal HAS-barrel domain; KEGG K06915 |
| sll0639 | 15 | 12 | 7, 12 | 417 | 0.74 | 0.63 | DHH phosphoesterase domain- |
| ssr1238, slr0743a | 15 | 11 | 11 |  | 0.71 | 1.04 | YlxR homologue |
| slr1660 | 17 | 18 | 18 | 192 | 0.91 | 1.41 | DUF3172 with N-terminal transmembrane domain |

Manual curation of the shortlist of 14 proteins that had passed the criteria of appropriate cluster assignment, unknown functional categorization, minimum phylogenetic conservation and minimum SVM score identified three proteins that are unlikely to be involved with nucleic acids due to known functions. These are Sll1515, the glutamine synthetase inactivating factor IF17 (Garcia-Dominguez et al., 1999), Slr0169, a CsoD1 ortholog that is part of the β-carboxysome, containing two bacterial microcompartment (BMC) domains (Kinney et al., 2011), and Sll1939, an FtsZ interacting Ftn6 homolog involved in cell division (Marbouty et al., 2009). Nevertheless, we kept them in the list because a small number of bifunctional proteins exist in bacteria that are active in metabolism and in controlling gene expression, including mechanisms in which RNA is bound (Commichau and Stülke, 2008).

Another four proteins might interact with nucleic acids but less so with RNA. These proteins are Ssl2874, a homolog of RemA, an essential regulator of biofilm formation in *Bacillus subtilis* (Winkelman et al., 2009, 2013), Sll0639, a DHH phosphoesterase domain-containing protein with similarities to a RecJ domain protein, nucleotidyltransferase/poly(A) polymerase or exopolyphosphatase-like enzyme, Slr1081 (DUF820) featuring a restriction endonuclease domain and Sll0284 (DUF87) containing an N-terminal HAS-barrel domain similar to archaeal proteins, such as DNA double-strand break repair helicase HerA. However, bacterial proteins containing similar domains seem to be largely undescribed.

Another four poorly characterized proteins were Slr0211, an acyl-CoA N-acyltransferase domain containing protein, Sll0319 (DUF3747), a periplasmic protein, Slr1660 (DUF3172) containing an N-terminal transmembrane domain and Ssl2064, a 75 amino acid DUF4160-containing protein.

UItimately, three candidates appeared most promising. The first was Sll1424, an undescribed protein with very conserved synteny to the transcriptional regulator NtcA (**Supplementary Figure S7A**). Furthermore, we noticed Ssr1238/Slr0743a, an 84 amino acid YlxR homologue with a DUF448 domain and a pronounced synteny in cyanobacteria, expressed from a gene between *nusA* encoding the transcription termination factor NusA and *infB* encoding the translation initiation factor InfB (**Supplementary Figure S7B**). While the order of adjacent genes *rimP-nusA-infB* is conserved even in *Salmonella* and *E. coli*, a gene corresponding to *ssr1238* is lacking in their genomes. The relationship of *ylxR* with *rimP-nusA-infB* has also been observed in recent analyses of cyanobacterial gene linkage networks (Labella et al., 2020b).

Finally, we identified Slr0287, a KH domain-containing 118 amino acid protein that shares 30% identical and 56% similar residues with the corresponding protein KhpA from *S. pneumonia*. KhpA forms together with the JAG/EloR-domain protein KhpB and RNA-binding complex in *S. pneumonia* (Zheng et al., 2017). Indeed, we noticed that with Slr1472, a KhpB homolog also exists in *Synechocystis*, and both proteins are widely conserved throughout the cyanobacterial phylum and even cluster together in the gradient (**Supplementary Figure S5**). The genes of both proteins are syntenic with genes encoding proteins involved in RNA metabolism and translation, such as *khpA/slr0287* with *rps16* (**Supplementary Figure S7C**) or *khpB/slr1472* with *rpl34, rnpA and slr1470* encoding a membrane protein and *slr1471* encoding a YidC/Oxa1 homolog (**Supplementary Figure S7D**). Slr1471/Oxa1 is essential for thylakoid biogenesis and the membrane integration of the reaction center precursor protein pD1 (Spence et al., 2004; Ossenbühl et al., 2006). The synteny of *rnpA, yidC/oxa1* and *khpB* as well as *khpA* and *rps16* is strikingly wide, extending even to gram-positive bacteria, such as *S. pneumonia* (not shown).

### Data accessibility and visualizations

To facilitate maximum usability of this resource, we have made the entire dataset available at https://sunshine.biologie.uni-freiburg.de/GradSeqExplorer/, providing multiple tools to focus on different fractions of the proteome or transcriptome, from groups of functionally related proteins and transcripts to the level of the individual protein or sRNA. Multiple filter options, a customizable graphical output, as well as a tabular output options provide extensive background information to the findings presented in this analysis.

## Discussion

### Global analysis of RNA-protein complexes in a photosynthetic cyanobacterium

Based on the interactions with different types of RBPs, five major classes of ncRNAs were differentiated that are common to many bacteria (Hör et al., 2018). These include the 6S RNA as a global modulator of RNA polymerase (RNAP) specificity and activity (Barrick et al., 2005; Wassarman and Storz, 2000), the Hfq-binding sRNAs (Møller et al., 2002; Zhang et al., 1998; Moll et al., 2003), the ProQ-binding sRNAs (Smirnov et al., 2016), the CsrB-like sRNAs that antagonize the translational repressor CsrA (Liu et al., 1997), and CRISPR RNAs (crRNAs), which form together with the CRISPR-associated (Cas) proteins ribonucleoprotein particles for defense (Hale et al., 2009; Brouns et al., 2008). Trans-acting sRNAs and their effects on gene expression have been well studied in *Escherichia coli* and *Salmonella enterica* (Hör et al., 2020c) but much less so in photosynthetic cyanobacteria. While 6S RNA and crRNAs exist in cyanobacteria, the affiliation of certain sRNAs to certain RBPs has remained unresolved.

Our Grad-seq analysis of *Synechocystis* provides the first global analysis of major stable RNA-protein complexes in a photosynthetic cyanobacterium. We anticipate that our data not only provide crucial knowledge for this particular model cyanobacterium but also constitute a valuable resource for other cyanobacteria, including the models *Anabaen*a (*Nostoc*) sp. PCC 7120 or *Synechococcus elongatus*.

*Synechocystis* as a cyanobacterial model deviates substantially from other bacteria investigated in previous Grad-seq studies (Smirnov et al., 2016; Hör et al., 2020b). One aspect is the magnitude of internal membrane systems and membrane-bound protein complexes. We showed that even macromolecular membrane-bound protein complexes remained largely intact during the process, and their in-gradient localization matched the pigment composition of the gradient in a meaningful way. Hence, the analysis provides a spatially resolved proteome dataset of high quality for the macromolecular complexes of a photosynthetic bacterium. In addition, the occurrence of subunits specific for certain subcomplexes or specialized functions provides information about the *in vivo* state and illustrates the dynamics in complex formation. Examples for such acclimation processes include the incorporation of PsaK2 into PSI as the high light-adapted form instead of the constitutive form, which incorporates mainly PsaK1. In our data, the occurrence of PsaK2 was restricted to disassembled and monomeric PSI, while PsaK1 occurred mainly in fully assembled monomeric and trimeric PSI, capturing the *in vivo* state of *Synechocystis* PSI and a multitude of other complexes under standard growth conditions.

Our data not only provide information about known protein-protein interactions but also provide clues about currently unknown interactions, such as the given examples of the PSII assembly factors Sll0933, Slr1768 and Slr2013, the PSI assembly factor VIPP1 homolog, the P_II_ interacting protein PipX or the membrane protein PamA with unexplained occurrences in the heavy fractions and outside the range of their known interaction partners, which illustrate the potential value of this dataset.

### Protein candidates for the interaction with sRNAs

To our surprise, the gradient profiles of the majority of RNAs diverged vastly from most detected proteins, locating generally in the heavier fractions and with largely similar sedimentation patterns, independent of transcript lengths. This fact is a strong indication of association with one or more major RNA chaperones or to large intact protein complexes involved in transcription or translation, such as RNAP, ribosomes and associated factors.

The colocalization of many known intact RNA-protein complexes in fractions 8 - 11 (cluster 11-15) involved in transcription, RNA metabolism and translation initiation shows the stability of such complexes, while the co-occurrence of a large number of RNAs illustrates that those complexes were still capturing the *in vivo* state of *Synechocystis* regulatory processes by binding RNAs. Furthermore, the fact that these fractions constituted a poor protein complexity but still harbored most of the well-studied factors involved in protein biosynthesis makes them interesting for the identification of novel candidates functionally linked to posttranscriptional regulation.

Another surprising observation was that the majority of detected transcripts were largely not associated with the previously mentioned complexes but mainly sedimented in fractions 6-7 (cluster 10), indicating the involvement of one or more RNA chaperones stably binding their target RNAs. The data permit for the first time the identification of candidate RNA-binding proteins involved in the formation of major RNA–protein complexes in this group of bacteria. Although *Synechocystis* expresses multiple sRNAs of crucial relevance for oxygenic photosynthesis and other important physiological processes (Klähn et al., 2015; Georg et al., 2017, 2014; de Porcellinis et al., 2016), homologs of CsrA, ProQ or Hfq do not exist in cyanobacteria or are not RNA binding (Schuergers et al., 2014). A prediction from our Grad-seq cluster analysis is that cyanobacteria may rather not utilize a general RBP analogous to Hfq or ProQ as characterized in enterobacteria, in which these two RBPs together are associated with more than 80% of all sRNAs (Holmqvist and Vogel, 2018). On the one hand, sRNAs in cyanobacteria may frequently function independent of a common RNA chaperone, or they may rely more upon specialized RBPs. In this respect, our data resemble the conclusions recently drawn for gram-positive bacteria based on the Grad-seq analysis of the human pathogen *S. pneumonia* (Hör et al., 2020b). We filtered out a short list of previously uncharacterized protein candidates for the interaction with sRNAs. Among them, we discovered a cyanobacterial homolog of the KhpA/B complex (Zheng et al., 2017). The nearly congruent peaks of the homologs Slr0287 and Slr1472 in gradient fractions 8–10 indicate that these proteins are part of a higher molecular weight complex, actually much larger than the complex reported for *S. pneumonia*, which sedimented in fractions 2 and 3.

### Limitations of the study

The here presented dataset is of high complexity indicated by the numbers of several hundreds up to more than 2,000 different proteins and high numbers of transcripts detected in the three replicates of every single fraction (**Supplementary Table S1**). This complexity is not solely a representation of the true biological complexity but in part resulting from technical variations which were taken into account by the chosen clustering approach. We detected that some subcomplexes likely became unstable during the procedure and dissociated from the respective main complex, such as the oxygen evolving complex of PSII. Similarly, PSI appears to have been almost completely monomerized (**Figure 5**), CP43 (PsbC) peaked in F4/F5, whereas the other PSII core complex subunits were found in F6/F7 (**Figure 5**), and also NDH-I subunits (**Supplementary Figure S4**) were distributed through fractions 3 to 6 of the gradient with different abundances, also indicative partial complex disruption. Hence, the possibility that some complexes were fully stable only when membrane bound, cannot be excluded and cannot be avoided during the solubilization of the membranes. Moreover, the high number of detected proteins together with a relatively low number of fractions limited the resolution, especially in the lower fractions F1 to F6 with high protein complexity. On the other hand, the resolution appeared better in the higher fractions F7 to F18, where most transcripts and large protein complexes were detected while the protein complexity was lower, making these fractions suitable for the detection of RNA:protein complexes. Together with the chosen clustering approach to divide the content of the dataset into groups of similarly sedimenting proteins and transcripts and the extended bioinformatic analysis, this has aided the identification of previously uncharacterized protein candidates likely involved in the metabolism of RNA and post-transcriptional regulation.

## Materials and Methods

### Cell cultivation, cell lysis, gradient preparation and fractionation

Triplicates of 400 mL of *Synechocystis* liquid cultures were cultivated in BG11 medium under constant shaking in standard growth conditions (30 °C, 50 µE) until reaching exponential phase (OD_750_ = 0.8). The cultures were harvested by centrifugation at 4000 g for 15 min at RT. After centrifugation, the cell pellets were resuspended in ice cold lysis buffer (20 mM Tris-HCl, pH 7.5; 150 mM KCl; 1 mM MgCl_2_; 1 mM DTT; 0.2 % [v/v] Triton X-100; RiboLock RNase Inhibitor; DNase I; cOmplete Protease Inhibitor cocktail) and mechanically lysed with glass beads at 4 °C (Precellys). From here on, the samples were kept at 4 °C during the whole procedure. Unbroken cells were removed by a centrifugation step for 5 min at 4000 g, and the supernatant was transferred to a fresh tube. The membrane proteins were solubilized by incubating the samples under shaking for 60 min in the dark with n-dodecyl β-D-maltoside (DDM) in a 3:1 ratio according to the protein content (1 % [w / v], 1 g / 100 mL initial culture). RNase inhibitor was added to minimize the risk of RNA degradation during the 1 h solubilization and the visual inspection of RNA preparations after fractionation showed no sign of substantial RNA degradation. Thylakoid membranes were removed by centrifugation at 30,000 g for 30 min, and the supernatant was applied on top of linear sucrose gradients. We modified the original Grad-seq protocol by using DDM instead of Triton X100. DDM is a mild detergent to allow the inclusion of membrane proteins as it is an efficient membrane solubilizer that is frequently used for the investigation of photosynthetic membrane complexes, both in cyanobacteria and in plants (Eshaghi et al., 2000; Golub et al., 2020; Barera et al., 2012; Ma et al., 2006). Moreover, based on the results of pilot experiments we generated the density gradients from sucrose instead of glycerol. The 10 - 40 % sucrose gradients were made using a gradient mixer with two solutions of lysis buffer, supplemented with either 10 % or 40 % sucrose [w / v].

Ultracentrifugation of the lysates was performed in a swing out rotor (Beckman SW40 Ti) for 16 h at 285,000 g. The fractionation of equal volumes (600 µL) was performed manually, and the fractions were split into samples of equal volumes for MS analysis (100 µL) and library preparation for RNA sequencing (360 µL). The protein complexity of the individual fractions were determined in pre-experiments which revealed low complexity of fractions 14 - 17. Therefore, fractions 14 and 15 (F1415), as well as fractions 16 and 17 (F1617) were pooled prior to downstream sample preparations out of economic considerations, resulting in a total of 16 samples per replicate for transcriptome and proteome analyses. Fraction 18 represents the gradient pellet.

### Spectroscopy

Absorption spectra were measured using a Specord® 250 Plus (Analytik Jena) spectrophotometer at room temperature and were normalized to equal fraction volumes.

### RNA sample preparation, generation of cDNA libraries and sequencing

The 360 µL fraction samples were spiked with a barcoded 197 bp long spike-in RNA mix ranging from 1 - 100 fmol (sum 200 fmol) and subjected to proteinase K treatment (1 % SDS, 10 mM Tris-HCl pH 7.5, proteinase K) for 30 min at 50 °C, followed by RNA extraction with PGTX (Pinto et al., 2009). After the first phase separation step, the aqueous phase was transferred to a fresh microcentrifuge tube and mixed 1:1 [v / v] with 100 % EtOH, and RNA was further extracted using the “RNA clean & concentrator” kit from Zymo according to the manufacturer’s instructions. The spike-in RNA mix was created by in vitro transcription (Mega-shortscript, Thermo Fisher) of PCR-amplified parts from the genomic DNA of a phiX174 bacteriophage genome (**Supplementary Table S4**), purified from a 10% urea-PAA gel (RNA PAGE Recovery, (Zymo). Determination of RNA concentration was performed using the Qubit RNA HS Assay kit (Thermo Fisher). For the control, equal volumes of purified RNA were separated on a 10% denaturing urea-PAA gel.

The library preparation and Illumina paired end sequencing (75 bp read length) were performed by vertis Biotechnologie AG. In short, RNA samples were fragmented using ultrasound (4 pulses of 30 s each at 4 °C), followed by 3’ end adapter ligation and first strand cDNA synthesis using M-MLV reverse transcriptase. The 5’ Illumina TrueSeq sequencing adapter was ligated to the 3’ end of the antisense from the purified first-strand cDNA, and the resulting cDNA was PCR amplified (11 cycles) and purified using the Agencourt AMPure XP kit (Beckman Coulter Genomics). For Illumina NextSeq sequencing, the samples were pooled, and the cDNA pool was eluted in the size range of 160 - 500 bp from a preparative agarose gel. The primers used for PCR amplification were designed for TruSeq sequencing according to the instructions of Illumina (**Supplementary Table S4**).

### Read mapping and normalization

The analysis of the raw reads from the RNA-sequencing was performed using the galaxy web platform. The workflow can be accessed and reproduced with the following link: https://usegalaxy.eu/u/mr559/w/-grad-seq-pipeline. In short, the read quality was checked at the beginning and after each step with FastQC. Adapter trimming was performed using Cutadapt (Martin, 2011), and the trimmed reads were mapped to the *Synechocystis* genome using segemehl (Otto et al., 2014) with default parameters and were assigned to annotated regions using FeatureCounts. Prior to normalization, all features with a maximum read count ≤ 5 across all fractions were filtered prior to downstream analysis. Fraction-wise normalization was performed via the spike-in RNA mix which was added in equal amounts to each fraction prior RNA sample preparation. The relative RNA amounts per transcript refer to the spike-in normalized reads across the gradient. The reproducibility between the replicates was checked by comparing the distribution of the Spearman correlation coefficient (calculated with ‘cor()’ function from R stats package) from the relative distribution of each RNA within a replicate with the same RNAs in the other replicates (**Supplementary Figure S1A**).

### Sample preparation for mass spectrometry-based proteome measurements

As an initial step, proteins were purified by acetone/methanol precipitation. Therefore, 100 µL of each of the initial 18 fractions were mixed with an ice-cold mixture of 800 µL acetone and 100 µL methanol. After incubation o.n. at −20 °C, precipitated proteins were subsequently collected by centrifugation at 1000 × g and 4 °C. Subsequently, protein pellets were washed twice with 1 mL cold 80 % (v / v) acetone (aq.). During the washing procedure, precipitated sucrose was removed almost completely, also in fractions with high sucrose content.

Dried protein pellets were consequently redissolved in 50 µL denaturation buffer (6 M urea, 2 M thiourea in 100 mM Tris/HCl, pH 7.5). Intrachain disulfide bonds were reduced with dithiothreitol and resulting thiol groups were alkylated with iodoacetamide as described before (Spät et al., 2015). Subsequently, proteins were predigested with endoprotease Lys-C for 3 h, further diluted with 200 µL 20 mM ammonium bicarbonate buffer, pH 8.0, and digested with trypsin o.n., applying a protease:protein ratio of 1:100 for both enzymes. The resulting peptide solutions were acidified to pH 2.5 with 10 % (v / v) trifluoroacetic acid (aq.) and purified following the stage tip protocol (Rappsilber et al., 2007).

For MS-based proteome measurements, a constant sample volume was analyzed for all fractions of each replicate to account for the different protein amounts between the fractions. Consequently, the measured protein yields ranged between a maximum of 1000 ng and <100 ng in fractions with the lowest protein concentrations (**Supplementary Table S5**). The relative protein amounts refer to the relative iBAQ intensities from equal fraction volumes across the gradient (equal volume). As a control, equal volumes of the initial gradient fractions were separated in 15% SDS-PAA gels and visualized by Coomassie staining.

### MS measurements

For nanoLC-MS/MS-based proteome measurements, purified samples were loaded onto an in-house produced C18 nanoHPLC column (20 cm PicoTip fused silica emitter with 75 µm inner diameter (New Objective, Woburn, USA), packed with 1.9 µm ReproSil-Pur C18-AQ resin (Dr Maisch, Ammerbuch, Germany)) on an EASY-nLC 1200 system (Thermo Scientific, Bremen, Germany). Peptides were separated using a 49 min segmented linear gradient (**Supplementary Table S6**), and eluting peptides were directly ionized through an on-line coupled electrospray ionization (ESI) source and analyzed on a Q Exactive HF mass spectrometer (Thermo Scientific, Bremen, Germany), operated in the positive-ion mode and data dependent acquisition. MS full scans were acquired with a mass-to-charge (m/z) range of 300 - 1650 at a resolution of 60,000. The 12 most intense multiple charged ions were selected for fragmentation by higher-energy collisional dissociation (HCD) and recorded in MS/MS scans with a resolution of 30,000. Dynamic exclusion of sequenced precursor ions was set to 30 s. Wash runs (**Supplementary Table S6**) were performed after each of the three samples.

### Proteomics data processing

Acquired raw MS data of all three replicates were processed using the MaxQuant software suite (version 1.5.2.8) (Cox and Mann, 2008). Each file was defined as an individual sample. The data processing was performed at default settings, applying intensity based absolute quantification (iBAQ) and the following search criteria: trypsin was defined as cleaving enzyme and a maximum of two missed cleavages was allowed. Carbamidomethylation of cysteine residues was set as a fixed modification and methionine oxidation and protein N-terminal acetylation were defined as variable modifications. MS/MS spectra were searched against a combined target-decoy database of *Synechocystis* downloaded from Cyanobase (http://genome.microbedb.jp/cyanobase; version 06/2014), and the sequences of newly discovered small proteins (Mitschke et al., 2011; Kopf et al., 2014; Baumgartner et al., 2016), representing a total of 3,681 protein sequences plus 245 common contaminants. To enable the identification of low abundant proteins which were not selected for MS/MS sequencing in any fraction, ancillary measurements were performed for the first replicate with a constant protein yield of 1000 ng per fraction. These supporting measurements were included in the data processing as a resource to promote overall protein identifications via the matching between runs option. Reported protein and peptide false discovery rates were each limited to 1%. The reproducibility between the replicates was checked by comparing the distribution of the Spearman correlation coefficient (calculated with ‘cor()’ function from R stats package) from the relative distribution of each protein within a replicate with the same protein in the other replicates (**Supplementary Figure S1B**). An additional feature to visualize constant protein amounts per fraction was implemented into the web application (constant protein amount). This proved to be useful for the detection of lowly abundant protein sub-complexes, which were within the gradient in close proximity of highly abundant other sub-complexes (e.g. PSI monomer and PSI trimer). The constant protein amount was estimated by determining the protein amount in equal fraction volumes divided by the total protein amount per fraction. The first replicate, which was also measured with a constant protein amount of 1000 ng per fraction served as reference for the quality of this estimation.

### Hierarchical clustering and module detection

Clustering and module (cluster) detection were performed using the dynamic tree cut algorithm implemented in the WGCNA R package (Langfelder and Horvath, 2008, 2012) from the standardized relative abundances (z-scores, calculated with ‘scale()’ function from R base package) of the normalized RNA-seq reads and IBAQ intensities. The unsupervised detection of 17 clusters was mainly dependent on the gradient sedimentation profiles of the proteins and transcripts from the input data and their variations between the replicates. The parameters for the hierarchical clustering method and the dynamic tree cut algorithm were adjusted according to the standard procedure of WGCNA (Zhang and Horvath, 2005). The Pearson correlation coefficients of the individual protein and transcript distribution to the assigned cluster means were calculated with the ‘cor()’ function from the R stats package.

### RNApred benchmarking for *Synechocystis*

The performance of RNApred was tested for *Synechocystis* proteins categorized into Gene Ontology groups “Nucleotide binding”, “DNA binding”, “RNA binding” or “No binding” if no other category was assigned. The majority of the “no binding” group did not reach an SVM score >0.49, while the majority of the “RNA binding” group reached an SVM score of at least 0.26 (**Supplementary Figure S6A**). The distribution of those groups barely overlapped, showing the validity of RNApred for *Synechocystis* proteins. Furthermore, it was tested against all ribosomal proteins (**Supplementary Figure S6B**) and selected known RNA-binding proteins (**Supplementary Figure S6C**). Notably, the cyanobacterial Hfq failed to fulfill the set criteria by far, matching previous observations of *Synechocystis* Hfq as not binding RNA (Schuergers et al., 2014), thus confirming this approach. The groups “Nucleotide binding” and “DNA binding” were distributed between those two other groups, with the majority of the “DNA binding” group below an SVM score of 1.07. Therefore, a conservative minimum SVM score of at least ≥ 0.5 should serve sufficient to remove most proteins without nucleic acid binding properties. The median SVM score of all orthologs in the genomes give in **Supplementary Table S3** was included to enhance the robustness of the approach, applying a minimum score of ≥ −0.2, which is the default threshold for RBPs in the webserver. The boxplots for the SVM scores were generated using the ‘geom_boxplot()’ function of the ggplot2 R package. The upper and lower hinges correspond to the 75^th^ and 25^th^ percentiles. The median is shown as black line within the boxes. The outliers are shown as black dots outside of the whisker ranges. The *Synechocystis* SVM scores are shown separately as red dots.

### Data availability

The datasets produced in this study are available in the following databases:

- FastQ files from RNA sequencing in SRA (BioProject accession number PRJNA608723): https://www.ncbi.nlm.nih.gov/sra/PRJNA608723
- Galaxy workflow used for raw read processing: https://usegalaxy.eu/u/mr559/w/-grad-seq-pipeline
- All code involving the bioinformatics workflow and the shiny app is available on Github (https://github.com/MatthiasRiediger/Analysis-of-a-photosynthetic-cyanobacterium-rich-in-internal-membrane-systems-via-Grad-seq.git)
- Mass spectrometry raw data deposited at the ProteomeXchange Consortium (http://proteomecentral.proteomexchange.org) via the PRIDE partner repository (Vizcaíno et al., 2013) under the identifier: PXD020025
- Data accessibility and visualization: https://sunshine.biologie.uni-freiburg.de/GradSeqExplorer/

## Supporting information

Supplemental Dataset 2 with Tables S1 to S6

## Acknowledgments

Funded by the Deutsche Forschungsgemeinschaft (DFG, German Research Foundation) - 322977937/GRK2344, the priority program SPP2141 (grant HE 2544/14-1) to WRH and through the “SCyCode” research group 397695561/FOR 2816 to WRH and BM.

## Author contributions

WRH and MR designed the research and performed the experiments and bioinformatics analyses. PS and BM carried out the MS and the related analyses. MR developed the bioinformatics analysis workflow and database. RB performed validation experiments and KV set up the hosting webserver. WRH and MR wrote the manuscript with input from all authors. WRH and BM supervised and funded the project.

## Supplemental information

**Supplementary Tables** (see separate Supplemental Dataset 2 for Tables S1 to S6)

**Table S1: Overview on the fraction complexity**. The table includes the total numbers of all measured proteins and transcripts per fraction.

**Table S2: Full Grad-seq dataset**. The table includes the relative abundances (rounded to four decimal places) of all detected proteins and transcripts in triplicates along the gradient.

**Table S3: All selected genomes used for the phylogenetic occurrence analysis based on domclust of the Microbial Genome Database (MBGD)**.

**Table S4: List of oligonucleotides**.

**Table S5: Total protein content per fraction measured by Bradford assay**.

**Table S6: nanoHPLC gradient used for MS-based proteome measurements**.

## Supplementary Figures

**Figure S1.**
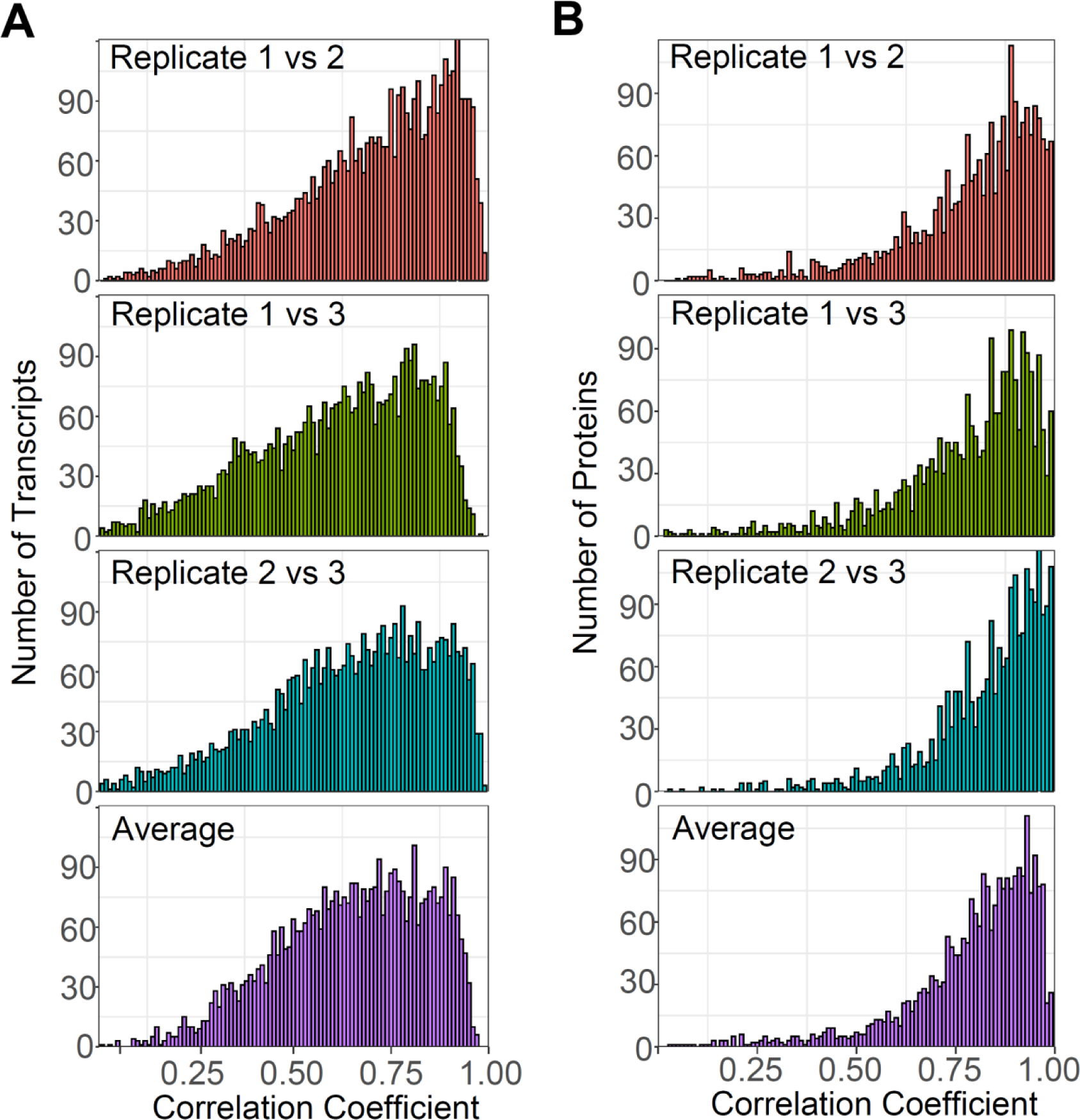
Histograms of the Spearman correlation coefficients from the comparison of gradient profiles between the replicates for **(A)** each detected RNA from the RNA sequencing and for **(B)** each detected protein from the MS analyses. This analysis supports **Figure 1**.

**Figure S2.**
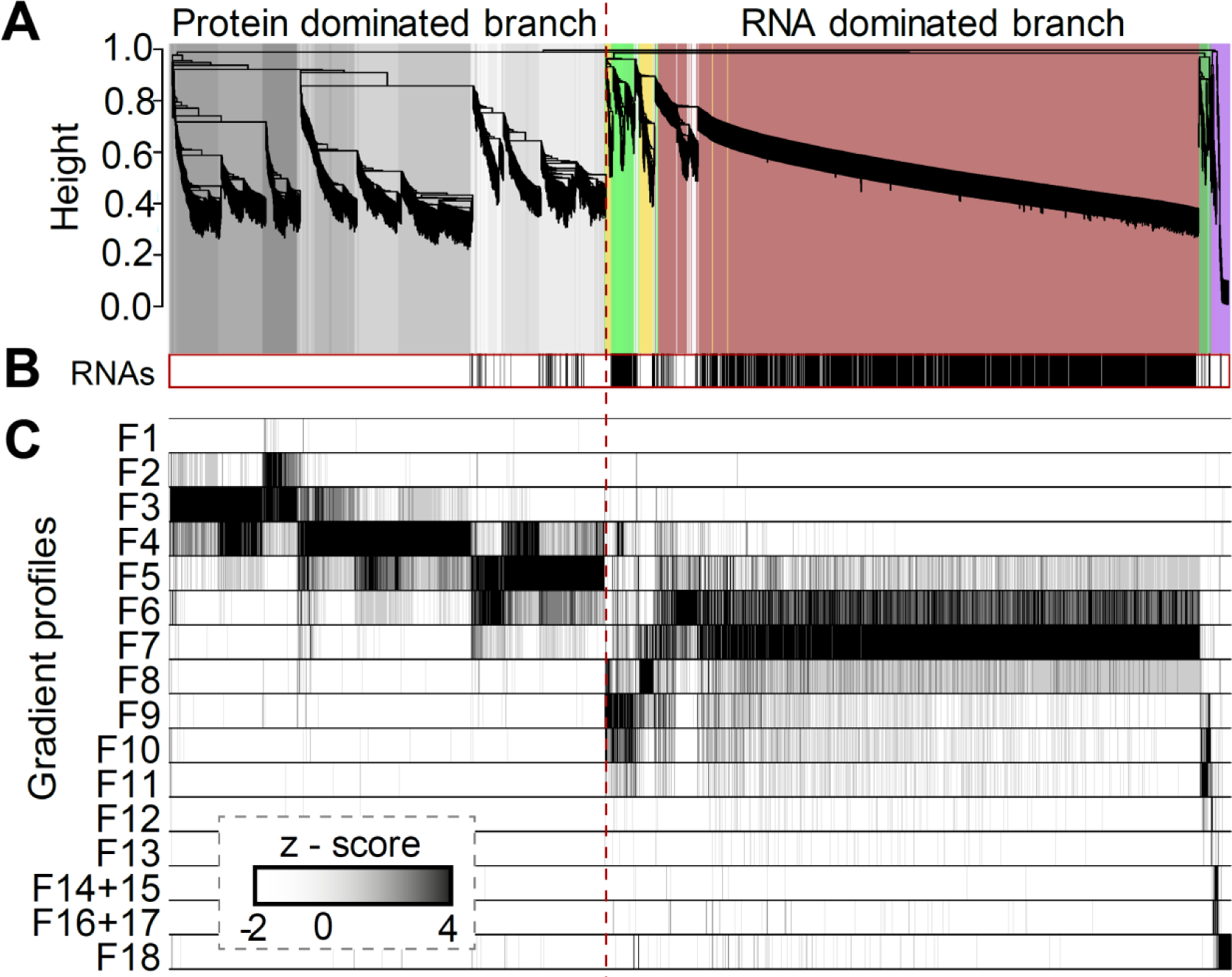
Distribution of 2394 proteins and >4000 different transcripts along a sucrose density gradient. **(A)** The hierarchical clustering approach using the standardized relative abundances of the normalized RNA-seq reads and MS intensities from the Grad-Seq experiment as input divides the dataset into 17 different clusters based on their distinct gradient profiles. The data bifurcate into two main branches, a protein-dominated and an RNA-dominated branch. The color code on the dendrogram represents the cluster affiliations. Gray-scale clusters are dominated by proteins that are distributed in the low-molecular-weight fractions (clusters 1-9). The colored clusters are dominated by the majority of RNAs and proteins that are distributed in the high-molecular-weight fractions (clusters 10-17). Details about the cluster composition are given in **Figure 2. (B)** Black strokes at the bottom of the branches of the dendrogram illustrate the RNA occurrences in these distinct clusters. **(C)** The heatmap illustrates the standardized average of the relative abundances (z-score) from three replicates for each detected protein and RNA alongside the fractions of the sucrose density gradient. This figure supports **Figure 2**.

**Figure S3.**
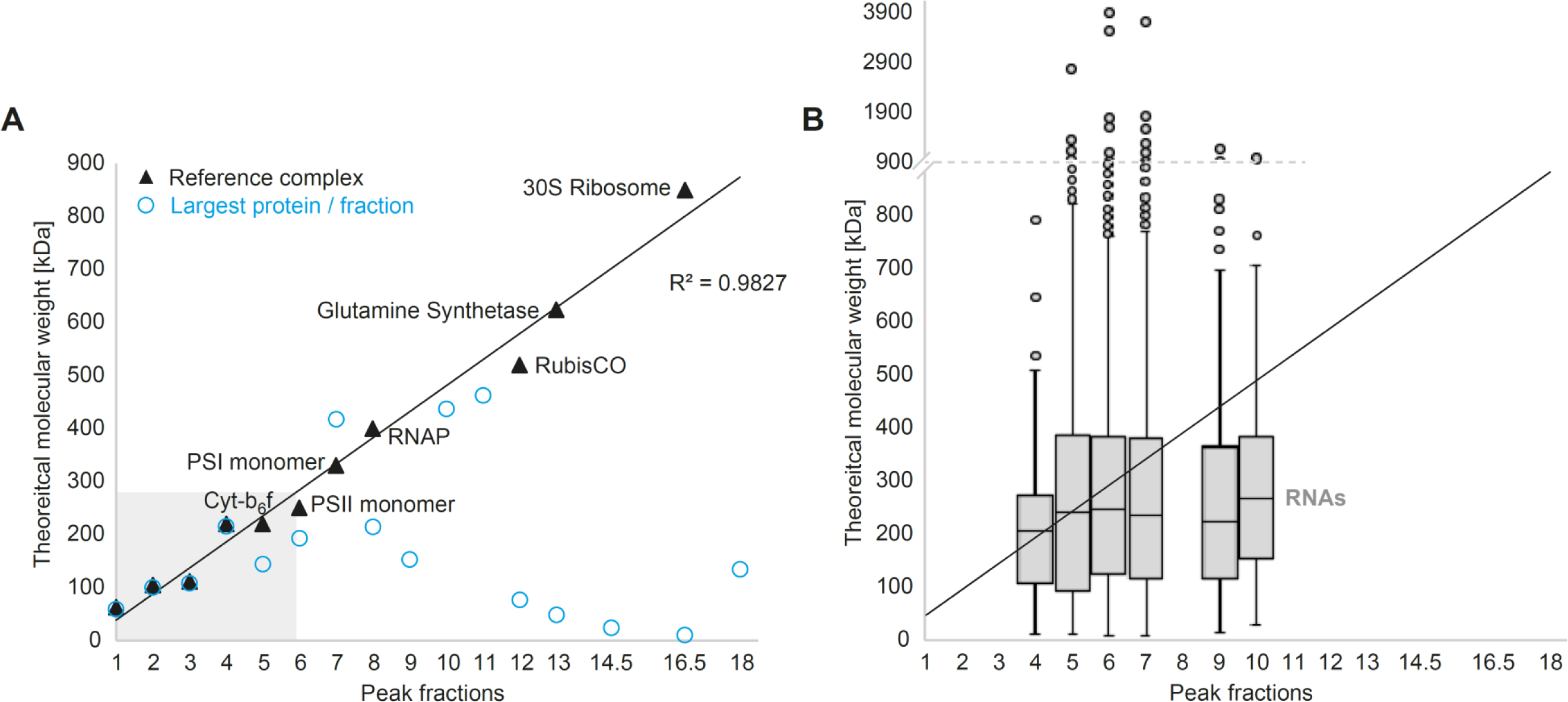
Sedimentation velocity in comparison to molecular weight. **(A)** Selected protein complexes (black triangles) illustrate the correlation between molecular weight and sedimentation velocity of proteins. The blue circles show the size of the largest single protein peaking in the respective fractions. The gray shaded area depicts the range of the lower molecular fractions harboring most soluble proteins and small-to medium-sized protein complexes. **(B)** Boxplots of the molecular weight of RNAs peaking in the respective fractions. This figure supports **Figure 2**.

**Figure S4.**
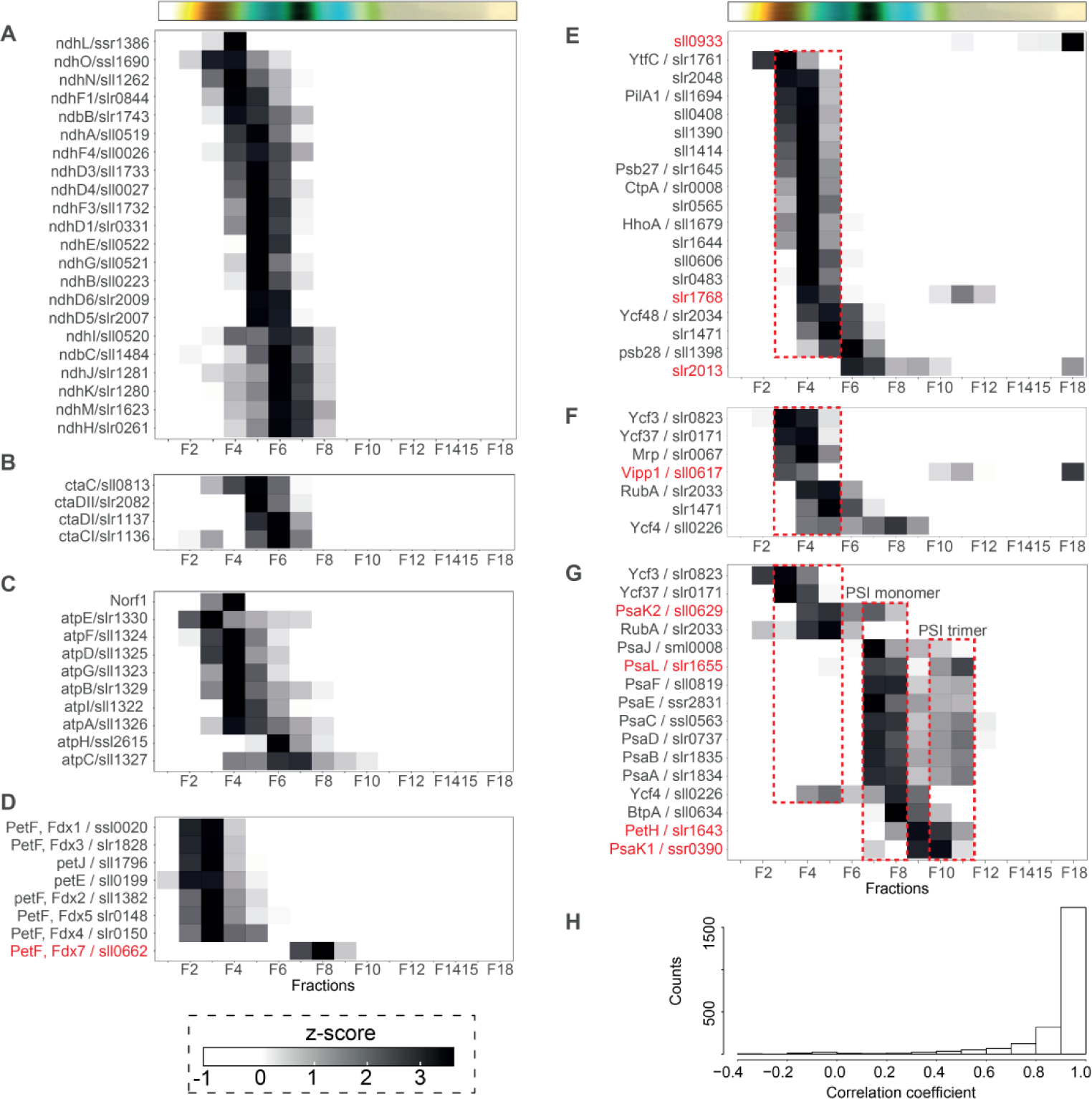
The heatmap illustrates the standardized relative abundances (z-score), giving the in gradient distribution of all detected **(A)** NADH dehydrogenase subunits, (**B**) cytochrome-c-oxidase complex subunits, **(C)** F0F1 ATP synthase subunits including Norf1, **(D)** the soluble electron carriers plastocyanin (PetJ), cytochrome c6 (PetE) and the ferredoxins, **(E)** PSII assembly factors reviewed in (Nickelsen and Rengstl, 2013) and **(F)** PSI assembly factors reviewed in (Yang et al., 2015). **(G)** In-gradient distribution of all detected photosystem I subunits normalized to constant protein amounts per fraction instead of constant volumes increases the visibility of the PSI trimer peak around fraction 10. Our measurements were based on equal volumes per fraction, which was chosen as it permits the comparison of proteomics with the RNA-seq data and avoids the pitfalls of the extremely different protein concentrations along the gradient fractions. **(H)** MS measurements for replicate 1 were performed with constant proteins as well as constant volumes per fraction. Therefore, the measurements of constant protein were compared to the measurements of constant volumes, which were afterwards corrected for equal protein amount by multiplication with the measured input concentrations, giving a good estimation for the quality of this approach. However, we only included this as an additional feature and did not perform extensive analyses based on the assumption of equal protein amounts per fraction. This figure extends the information given in **Figure 5**.

**Figure S5.**
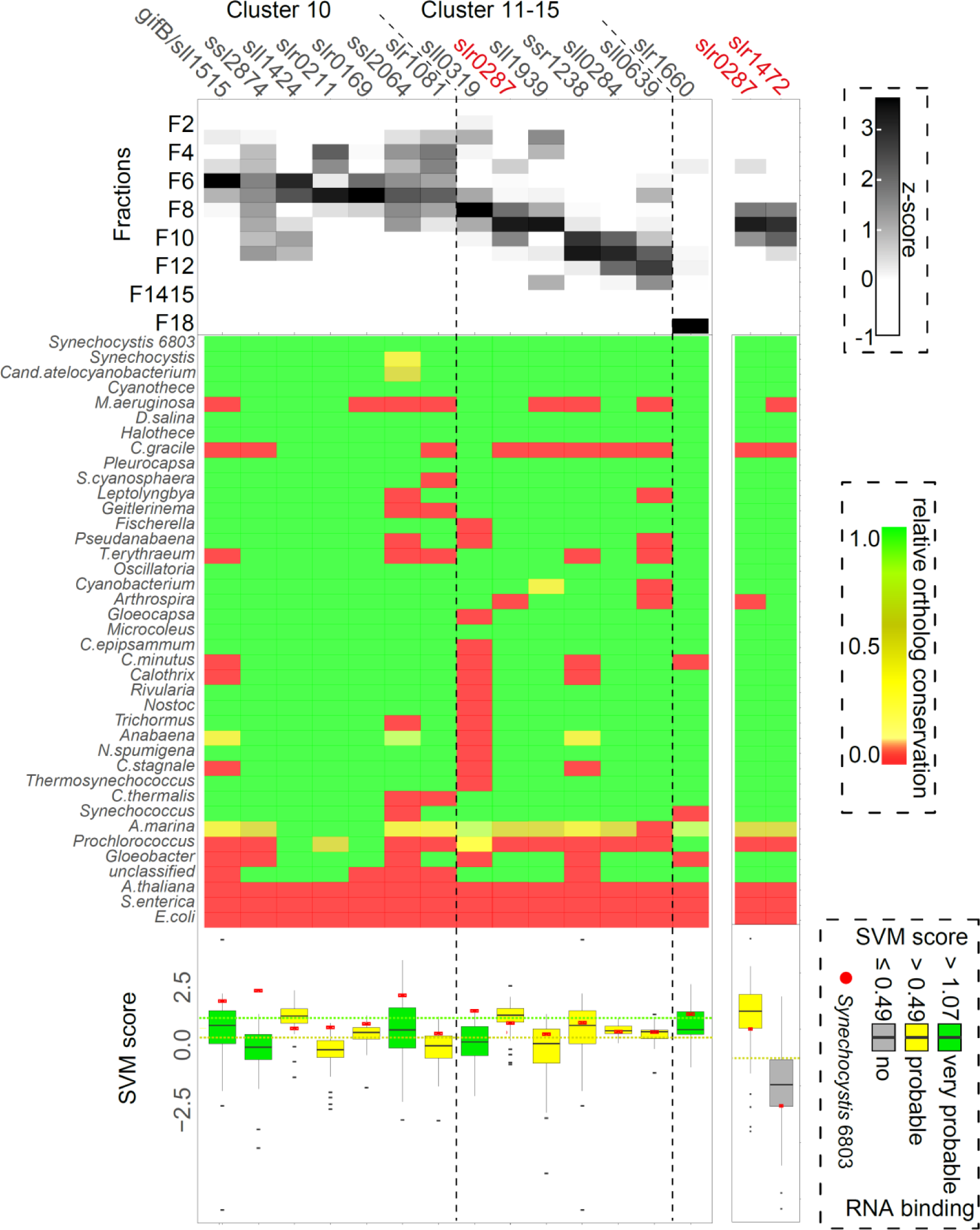
Cluster protein composition and selection process of putative cyanobacterial RNA binding proteins. All 14 selected RNA-binding protein candidates within the dataset were selected based on the previously mentioned prerequisites and additionally filter criteria according to high conservation among the selected cyanobacterial genomes (**Table S3**, at least 50%) as well as a good SVM score classification for RNA-binding proteins (SVM score ≥ 0.5 and median SVM score of all orthologs ≥ −0.2, further explanation in **Figure S6**). This figure relates to **Table 1**.

**Figure S6.**
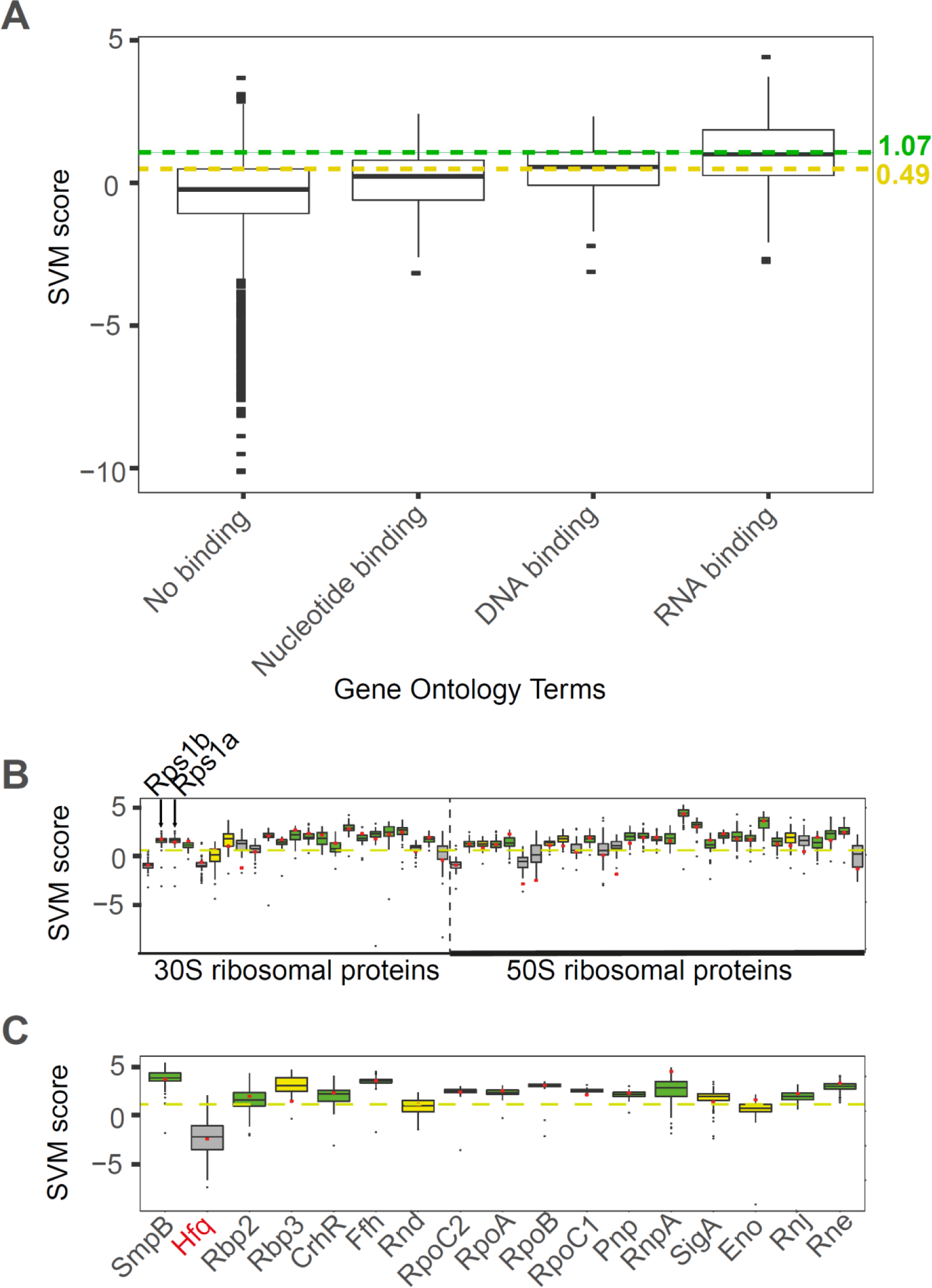
Comparison of determined SVM scores (Kumar et al., 2011) for proteins of *Synechocystis* with **(A)** Gene Ontology Terms (GO Terms) assigned either to “Nucleotide binding”, “DNA binding”, “RNA binding” or “No binding”. The limit of the upper quartile for “No binding” (0.49) was used as the threshold for the assignment of proteins into the yellow (probable RNA binding) group, whereas the limit of the upper quartile for “DNA binding” (1.07) was used as the threshold for the assignment of proteins into the green (very probable RNA binding) group. **(B)** Going into more detail, all Synechocystis ribosomal proteins (red dots) and their orthologs (boxplot) were tested for this approach, giving largely satisfying results, as well as selected **(C)** known RNA-binding proteins (red dots) and their orthologs (boxplot). The exception of Hfq not meeting the set SVM score criteria for cyanobacteria, and in particular for *Synechocystis*, once more illustrates its altered function as a non-RNA chaperone in cyanobacteria. This figure relates to **Table 1**.

**Figure S7.**
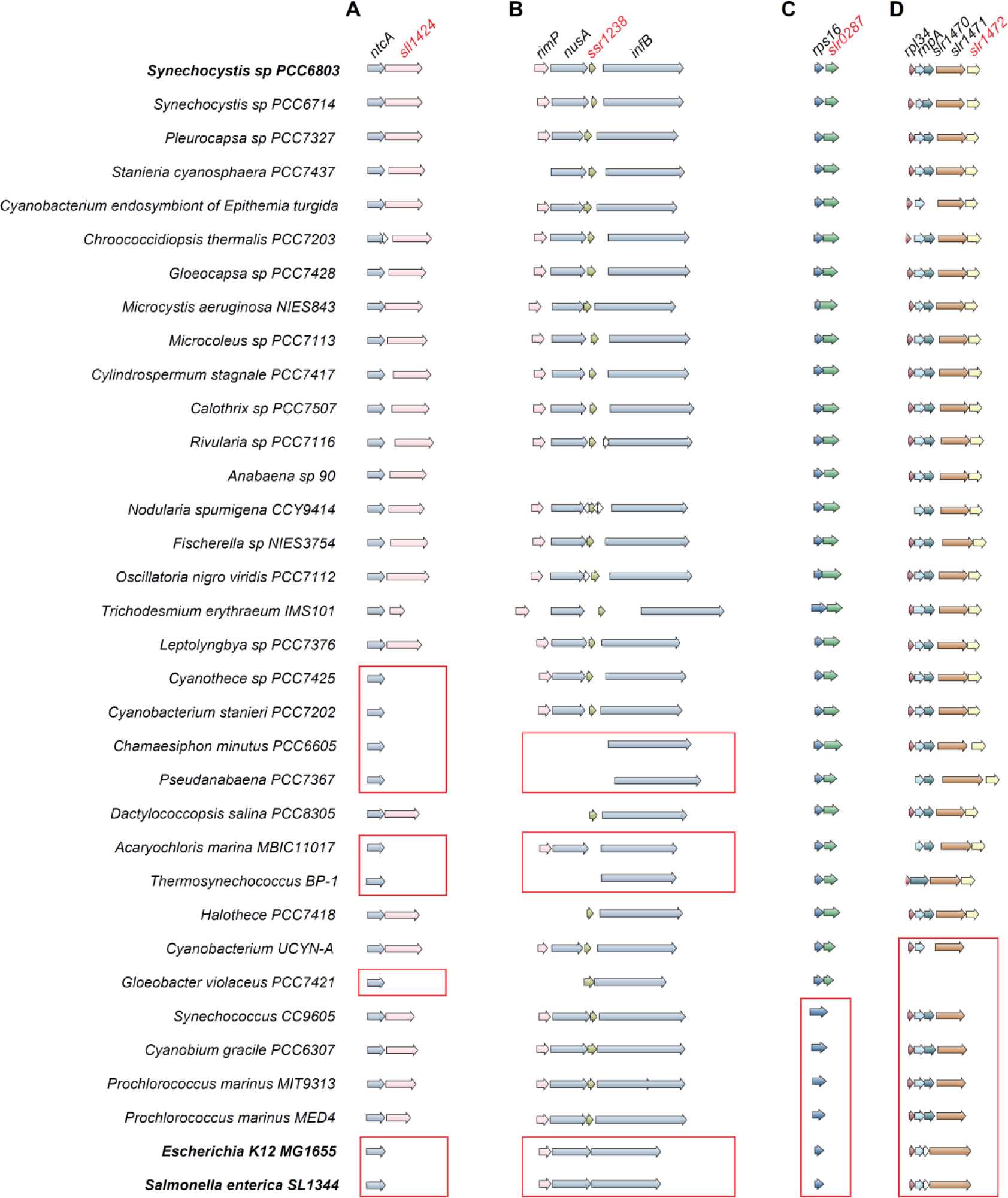
Conserved synteny in the cyanobacterial phylum of putative protein candidates involved in RNA metabolism. The synteny of **(A)** *ntcA* and *sll1424*, **(B)** *rimP, nusA, ssr1238* and *infB*, **(C)** *rps16* and *slr0287* and **(D)** *rpl34, rnpA, slr1470, slr1471* and *slr1472* is shown. The synteny was determined using the SyntTax web server (Oberto, 2013). This figure relates to **Table 1** and **supplementary Table S3**.

## References

Athukoralage, J.S., McQuarrie, S., Grüschow, S., Graham, S., Gloster, T.M., and White, M.F. (2020). Tetramerisation of the CRISPR ring nuclease Crn3/Csx3 facilitates cyclic oligoadenylate cleavage. eLife 9.

Baers, L.L., Breckels, L.M., Mills, L.A., Gatto, L., Deery, M.J., Stevens, T.J., Howe, C.J., Lilley, K.S., and Lea-Smith, D.J. (2019). Proteome mapping of a cyanobacterium reveals distinct compartment organization and cell-dispersed metabolism. Plant Physiol. 181: 1721–1738.

Bandyra, K.J. and Luisi, B.F. (2018). RNase E and the high-fidelity orchestration of RNA metabolism. Microbiol. Spectr. 6.

Barera, S., Pagliano, C., Pape, T., Saracco, G., and Barber, J. (2012). Characterization of PSII-LHCII supercomplexes isolated from pea thylakoid membrane by one-step treatment with α-and β-dodecyl-D-maltoside. Philos. Trans. R. Soc. Lond. B. Biol. Sci. 367: 3389–3399.

Barrick, J.E., Sudarsan, N., Weinberg, Z., Ruzzo, W.L., and Breaker, R.R. (2005). 6S RNA is a widespread regulator of eubacterial RNA polymerase that resembles an open promoter. RNA N. Y. N 11: 774–784.

Baumgartner, D., Kopf, M., Klähn, S., Steglich, C., and Hess, W.R. (2016). Small proteins in cyanobacteria provide a paradigm for the functional analysis of the bacterial micro-proteome. BMC Microbiol. 16: 285.

Behler, J., Sharma, K., Reimann, V., Wilde, A., Urlaub, H., and Hess, W.R. (2018). The host-encoded RNase E endonuclease as the crRNA maturation enzyme in a CRISPR–Cas subtype III-Bv system. Nat. Microbiol. 3: 367–377.

Bøggild, A., Overgaard, M., Valentin-Hansen, P., and Brodersen, D.E. (2009). Cyanobacteria contain a structural homologue of the Hfq protein with altered RNA-binding properties. FEBS J. 276: 3904–3915.

Brouns, S.J.J., Jore, M.M., Lundgren, M., Westra, E.R., Slijkhuis, R.J.H., Snijders, A.P.L., Dickman, M.J., Makarova, K.S., Koonin, E.V., and van der Oost, J. (2008). Small CRISPR RNAs guide antiviral defense in prokaryotes. Science 321: 960–964.

Cantos, R., Labella, J.I., Espinosa, J., and Contreras, A. (2019). The nitrogen regulator PipX acts in cis to prevent operon polarity. Environ. Microbiol. Rep. 11: 495–507.

Cech, T.R. and Steitz, J.A. (2014). The noncoding RNA revolution-trashing old rules to forge new ones. Cell 157: 77–94.

Chitnis, V.P. and Chitnis, P.R. (1993). PsaL subunit is required for the formation of photosystem I trimers in the cyanobacterium Synechocystis sp. PCC 6803. FEBS Lett. 336: 330–334.

Commichau, F.M. and Stülke, J. (2008). Trigger enzymes: bifunctional proteins active in metabolism and in controlling gene expression. Mol. Microbiol. 67: 692–702.

Cox, J. and Mann, M. (2008). MaxQuant enables high peptide identification rates, individualized p.p.b.-range mass accuracies and proteome-wide protein quantification. Nat. Biotechnol. 26: 1367–1372.

Dienst, D., Dühring, U., Möllenkopf, H.-J., Vogel, J., Golecki, J., Hess, W.R., and Wilde, A. (2008). The cyanobacterial homologue of the RNA chaperone Hfq is essential for motility of Synechocystis sp. PCC 6803. Microbiology 154: 3134–3143.

Dühring, U., Ossenbühl, F., and Wilde, A. (2007). Late assembly steps and dynamics of the cyanobacterial photosystem I. J. Biol. Chem. 282: 10915–10921.

Eshaghi, S., Turcsányi, E., Vass, I., Nugent, J., Andersson, B., and Barber, J. (2000). Functional characterization of the PS II-LHC II supercomplex isolated by a direct method from spinach thylakoid membranes. Photosynth. Res. 64: 179–187.

Forcada-Nadal, A., Llácer, J.L., Contreras, A., Marco-Marín, C., and Rubio, V. (2018). The PII-NAGK-PipX-NtcA regulatory axis of cyanobacteria: A tale of changing partners, allosteric effectors and non-covalent interactions. Front. Mol. Biosci. 5: 91.

Fujimori, T., Hihara, Y., and Sonoike, K. (2005). PsaK2 subunit in photosystem I is involved in state transition under high light condition in the cyanobacterium Synechocystis sp. PCC 6803. J. Biol. Chem. 280: 22191–22197.

Gao, F., Zhao, J., Chen, L., Battchikova, N., Ran, Z., Aro, E.-M., Ogawa, T., and Ma, W. (2016). The NDH-1L-PSI supercomplex is important for efficient cyclic electron transport in cyanobacteria. Plant Physiol. 172: 1451–1464.

Garcia-Dominguez, M., Reyes, J.C., and Florencio, F.J. (1999). Glutamine synthetase inactivation by protein-protein interaction. Proc. Natl. Acad. Sci. 96: 7161–7166.

Garcia-Doval, C., Schwede, F., Berk, C., Rostøl, J.T., Niewoehner, O., Tejero, O., Hall, J., Marraffini, L.A., and Jinek, M. (2020). Activation and self-inactivation mechanisms of the cyclic oligoadenylate-dependent CRISPR ribonuclease Csm6. Nat. Commun. 11: 1596.

Georg, J. et al. (2017). Acclimation of oxygenic photosynthesis to iron starvation is controlled by the sRNA IsaR1. Curr. Biol. 27: 1425-1436.e7.

Georg, J., Dienst, D., Schurgers, N., Wallner, T., Kopp, D., Stazic, D., Kuchmina, E., Klahn, S., Lokstein, H., Hess, W.R., and Wilde, A. (2014). The small regulatory RNA SyR1/PsrR1 controls photosynthetic functions in cyanobacteria. Plant Cell 26: 3661–3679.

Gerovac, M., Mouali, Y.E., Kuper, J., Kisker, C., Barquist, L., and Vogel, J. (2020). Global discovery of bacterial RNA-binding proteins by RNase-sensitive gradient profiles reports a new FinO domain protein. RNA 26: 1448–1463.

Golub, M., Hussein, R., Ibrahim, M., Hecht, M., Wieland, D.C.F., Martel, A., Machado, B., Zouni, A., and Pieper, J. (2020). Solution structure of the detergent-photosystem II core complex investigated by small angle scattering techniques. J. Phys. Chem. B 124: 8583–8592.

Gunnelius, L., Hakkila, K., Kurkela, J., Wada, H., Tyystjärvi, E., and Tyystjärvi, T. (2014). The omega subunit of the RNA polymerase core directs transcription efficiency in cyanobacteria. Nucleic Acids Res. 42: 4606–4614.

Hagemann, M. and Hess, W.R. (2018). Systems and synthetic biology for the biotechnological application of cyanobacteria. Curr Opin Biotechnol 49: 94–99.

Hale, C.R., Zhao, P., Olson, S., Duff, M.O., Graveley, B.R., Wells, L., Terns, R.M., and Terns, M.P. (2009). RNA-guided RNA cleavage by a CRISPR RNA-Cas protein complex. Cell 139: 945–956.

Han, W., Stella, S., Zhang, Y., Guo, T., Sulek, K., Peng-Lundgren, L., Montoya, G., and She, Q. (2018). A Type III-B Cmr effector complex catalyzes the synthesis of cyclic oligoadenylate second messengers by cooperative substrate binding. Nucleic Acids Res. 46: 10319–10330.

He, Z. and Mi, H. (2016). Functional characterization of the subunits N, H, J, and O of the NAD(P)H dehydrogenase complexes in Synechocystis sp. strain PCC 6803. Plant Physiol. 171: 1320–1332.

Heilmann, B., Hakkila, K., Georg, J., Tyystjärvi, T., Hess, W.R., Axmann, I.M., and Dienst, D. (2017). 6S RNA plays a role in recovery from nitrogen depletion in Synechocystis sp. PCC 6803. BMC Microbiol. 17: 229.

Holmqvist, E. and Vogel, J. (2018). RNA-binding proteins in bacteria. Nat. Rev. Microbiol. 16: 601–615.

Hör, J., Di Giorgio, S., Gerovac, M., Venturini, E., Förstner, K.U., and Vogel, J. (2020a). Grad-seq shines light on unrecognized RNA and protein complexes in the model bacterium Escherichia coli. Nucleic Acids Res. 48: 9301–9319.

Hör, J., Garriss, G., Di Giorgio, S., Hack, L.-M., Vanselow, J.T., Förstner, K.U., Schlosser, A., Henriques-Normark, B., and Vogel, J. (2020b). Grad-seq in a Gram-positive bacterium reveals exonucleolytic sRNA activation in competence control. EMBO J.: e103852.

Hör, J., Gorski, S.A., and Vogel, J. (2018). Bacterial RNA biology on a genome scale. Mol. Cell 70: 785–799.

Hör, J., Matera, G., Vogel, J., Gottesman, S., and Storz, G. (2020c). Trans-acting small RNAs and their effects on gene expression in Escherichia coli and Salmonella enterica. EcoSal Plus 9: 10.1128.

Hrle, A., Maier, L.-K., Sharma, K., Ebert, J., Basquin, C., Urlaub, H., Marchfelder, A., and Conti, E. (2014). Structural analyses of the CRISPR protein Csc2 reveal the RNA-binding interface of the type I-D Cas7 family. RNA Biol. 11: 1072–1082.

Ishikawa, Y., Schröder, W.P., and Funk, C. (2005). Functional analysis of the PsbP-like protein (sll1418) in Synechocystis sp. PCC 6803. Photosynth. Res. 84: 257–262.

Iyer, L.M., Koonin, E.V., and Aravind, L. (2004). Evolution of bacterial RNA polymerase: implications for large-scale bacterial phylogeny, domain accretion, and horizontal gene transfer. Gene 335: 73–88.

Kanehisa, M. (2019). Toward understanding the origin and evolution of cellular organisms. Protein Sci. 28: 1947–1951.

Kanehisa, M. and Goto, S. (2000). KEGG: kyoto encyclopedia of genes and genomes. Nucleic Acids Res. 28: 27–30.

Kanehisa, M., Sato, Y., Furumichi, M., Morishima, K., and Tanabe, M. (2019). New approach for understanding genome variations in KEGG. Nucleic Acids Res. 47: D590–D595.

Kazlauskiene, M., Kostiuk, G., Venclovas, Č., Tamulaitis, G., and Siksnys, V. (2017). A cyclic oligonucleotide signaling pathway in type III CRISPR-Cas systems. Science 357: 605–609.

Kieper, S.N., Almendros, C., Behler, J., McKenzie, R.E., Nóbrega, F.L., Haagsma, A.C., Vink, J.N.A., Hess, W.R., and Brouns, S.J.J. (2018). Cas4 facilitates PAM-compatible spacer selection during CRISPR adaptation. Cell Rep. 22: 3377–338.

Kinney, J.N., Axen, S.D., and Kerfeld, C.A. (2011). Comparative analysis of carboxysome shell proteins. Photosynth. Res. 109: 21–32.

Klähn, S., Bolay, P., Wright, P.R., Atilho, R.M., Brewer, K.I., Hagemann, M., Breaker, R.R., and Hess, W.R. (2018). A glutamine riboswitch is a key element for the regulation of glutamine synthetase in cyanobacteria. Nucleic Acids Res. 46: 10082–10094.

Klähn, S., Schaal, C., Georg, J., Baumgartner, D., Knippen, G., Hagemann, M., Muro-Pastor, A.M., and Hess, W.R. (2015). The sRNA NsiR4 is involved in nitrogen assimilation control in cyanobacteria by targeting glutamine synthetase inactivating factor IF7. Proc. Natl. Acad. Sci. 112: E6243–E6252.

Kondo, K., Geng, X.X., Katayama, M., and Ikeuchi, M. (2005). Distinct roles of CpcG1 and CpcG2 in phycobilisome assembly in the cyanobacterium Synechocystis sp. PCC 6803. Photosynth. Res. 84: 269–273.

Kondo, K., Mullineaux, C.W., and Ikeuchi, M. (2009). Distinct roles of CpcG1-phycobilisome and CpcG2-phycobilisome in state transitions in a cyanobacterium Synechocystis sp. PCC 6803. Photosynth. Res. 99: 217–225.

Kondo, K., Ochiai, Y., Katayama, M., and Ikeuchi, M. (2007). The membrane-associated CpcG2-phycobilisome in Synechocystis: A new photosystem I antenna. Plant Physiol. 144: 1200–1210.

Kopf, M. and Hess, W.R. (2015). Regulatory RNAs in photosynthetic cyanobacteria. FEMS Microbiol. Rev. 39: 301–315.

Kopf, M., Klähn, S., Scholz, I., Matthiessen, J.K.F., Hess, W.R., and Voß, B. (2014). Comparative analysis of the primary transcriptome of Synechocystis sp. PCC 6803. DNA Res. 21: 527–539.

Kopfmann, S., Roesch, S.K., and Hess, W.R. (2016). Type II toxin-antitoxin systems in the unicellular cyanobacterium Synechocystis sp. PCC 6803. Toxins 8: 228.1-228.23.

Kumar, M., Gromiha, M.M., and Raghava, G.P.S. (2011). SVM based prediction of RNA-binding proteins using binding residues and evolutionary information. J. Mol. Recognit. 24: 303–313.

Labella, J.I., Cantos, R., Salinas, P., Espinosa, J., and Contreras, A. (2020a). Distinctive features of PipX, a unique signaling protein of Cyanobacteria. Life 10: 79.

Labella, J.I., Llop, A., and Contreras, A. (2020b). The default cyanobacterial linked genome: an interactive platform based on cyanobacterial linkage networks to assist functional genomics. FEBS Lett. 594: 1661–1674.

Labella, J.I., Obrebska, A., Espinosa, J., Salinas, P., Forcada-Nadal, A., Tremiño, L., Rubio, V., and Contreras, A. (2016). Expanding the cyanobacterial nitrogen regulatory network: The GntR-like regulator PlmA interacts with the PII-PipX complex. Front. Microbiol. 7: 1677.

Langfelder, P. and Horvath, S. (2012). Fast R functions for robust correlations and hierarchical clustering. J. Stat. Softw. 46: i11.

Langfelder, P. and Horvath, S. (2008). WGCNA: an R package for weighted correlation network analysis. BMC Bioinformatics 9: 559.

Liu, H., Weisz, D.A., Zhang, M.M., Cheng, M., Zhang, B., Zhang, H., Gerstenecker, G.S., Pakrasi, H.B., Gross, M.L., and Blankenship, R.E. (2019). Phycobilisomes harbor FNR L in Cyanobacteria. mBio 10: e00669-19, /mbio/10/2/mBio.00669-19.atom.

Liu, M.Y., Gui, G., Wei, B., Preston, J.F., Oakford, L., Yüksel, U., Giedroc, D.P., and Romeo, T. (1997). The RNA molecule CsrB binds to the global regulatory protein CsrA and antagonizes its activity in Escherichia coli. J. Biol. Chem. 272: 17502–17510.

Llacer, J.L., Espinosa, J., Castells, M.A., Contreras, A., Forchhammer, K., and Rubio, V. (2010). Structural basis for the regulation of NtcA-dependent transcription by proteins PipX and PII. Proc. Natl. Acad. Sci. 107: 15397–15402.

Ma, W., Deng, Y., Ogawa, T., and Mi, H. (2006). Active NDH-1 complexes from the cyanobacterium Synechocystis sp. strain PCC 6803. Plant Cell Physiol. 47: 1432–1436.

Mahbub, M. et al. (2020). mRNA localization, reaction centre biogenesis and thylakoid membrane targeting in cyanobacteria. Nat. Plants 6: 1179–1191.

Makarova, K.S. et al. (2020). Evolutionary classification of CRISPR-Cas systems: a burst of class 2 and derived variants. Nat. Rev. Microbiol. 18: 67–83.

Makarova, K.S., Anantharaman, V., Grishin, N.V., Koonin, E.V., and Aravind, L. (2014). CARF and WYL domains: ligand-binding regulators of prokaryotic defense systems. Front. Genet. 5: 102.

Marbouty, M., Saguez, C., Cassier-Chauvat, C., and Chauvat, F. (2009). Characterization of the FtsZ-interacting septal proteins SepF and Ftn6 in the spherical-celled cyanobacterium Synechocystis strain PCC 6803. J. Bacteriol. 191: 6178–6185.

Martin, M. (2011). Cutadapt removes adapter sequences from high-throughput sequencing reads. EMBnet.journal 17: 10.

Melamed, S., Adams, P.P., Zhang, A., Zhang, H., and Storz, G. (2020). RNA-RNA interactomes of ProQ and Hfq reveal overlapping and competing roles. Mol. Cell 77: 411-425.e7.

Mitschke, J., Georg, J., Scholz, I., Sharma, C.M., Dienst, D., Bantscheff, J., Voß, B., Steglich, C., Wilde, A., Vogel, J., and Hess, W.R. (2011). An experimentally anchored map of transcriptional start sites in the model cyanobacterium Synechocystis sp. PCC6803. Proc. Natl. Acad. Sci. 108: 2124–2129.

Molina, R., Stella, S., Feng, M., Sofos, N., Jauniskis, V., Pozdnyakova, I., López-Méndez, B., She, Q., and Montoya, G. (2019). Structure of Csx1-cOA4 complex reveals the basis of RNA decay in Type III-B CRISPR-Cas. Nat. Commun. 10: 4302.

Moll, I., Leitsch, D., Steinhauser, T., and Bläsi, U. (2003). RNA chaperone activity of the Sm-like Hfq protein. EMBO Rep. 4: 284–289.

Møller, T., Franch, T., Højrup, P., Keene, D.R., Bächinger, H.P., Brennan, R.G., and Valentin-Hansen, P. (2002). Hfq: a bacterial Sm-like protein that mediates RNA-RNA interaction. Mol. Cell 9: 23–30.

Morris, K.V. and Mattick, J.S. (2014). The rise of regulatory RNA. Nat. Rev. Genet. 15: 423–437.

Mutsuda, M. and Sugiura, M. (2006). Translation initiation of cyanobacterial rbcS mRNAs requires the 38-kDa ribosomal protein S1 but not the Shine-Dalgarno sequence: development of a cyanobacterial in vitro translation system. J. Biol. Chem. 281: 38314–38321.

Nakagawa, S., Niimura, Y., Miura, K., and Gojobori, T. (2010). Dynamic evolution of translation initiation mechanisms in prokaryotes. Proc. Natl. Acad. Sci. U. S. A. 107: 6382–6387.

Nickelsen, J. and Rengstl, B. (2013). Photosystem II assembly: from cyanobacteria to plants. Annu. Rev. Plant Biol. 64: 609–635.

Niewoehner, O., Garcia-Doval, C., Rostøl, J.T., Berk, C., Schwede, F., Bigler, L., Hall, J., Marraffini, L.A., and Jinek, M. (2017). Type III CRISPR-Cas systems produce cyclic oligoadenylate second messengers. Nature 548: 543–548.

Ning, D., Zhao, W., and Qian, Y. (2013). [A hypothetical gene pair, ssl2138 and sll11092, constitutes a functional TA system on the chromosome of Synechocystis PCC 6803]. Wei Sheng Wu Xue Bao 53: 1043–1049.

Osanai, T., Sato, S., Tabata, S., and Tanaka, K. (2005). Identification of PamA as a PII-binding membrane protein important in nitrogen-related and sugar-catabolic gene expression in Synechocystis sp. PCC 6803. J. Biol. Chem. 280: 34684–34690.

Ossenbühl, F., Inaba-Sulpice, M., Meurer, J., Soll, J., and Eichacker, L.A. (2006). The Synechocystis sp PCC 6803 Oxa1 homolog is essential for membrane integration of reaction center precursor protein pD1. Plant Cell 18: 2236–2246.

Otto, C., Stadler, P.F., and Hoffmann, S. (2014). Lacking alignments? The next-generation sequencing mapper segemehl revisited. Bioinforma. Oxf. Engl. 30: 1837–1843.

Pinto, F., Thapper, A., Sontheim, W., and Lindblad, P. (2009). Analysis of current and alternative phenol based RNA extraction methodologies for cyanobacteria. BMC Mol. Biol. 10: 79.

de Porcellinis, A.J., Klähn, S., Rosgaard, L., Kirsch, R., Gutekunst, K., Georg, J., Hess, W.R., and Sakuragi, Y. (2016). The non-coding RNA Ncr0700/PmgR1 is required for photomixotrophic growth and the regulation of glycogen accumulation in the cyanobacterium Synechocystis sp. PCC 6803. Plant Cell Physiol. 57: 2091–2103.

Quendera, A.P., Seixas, A.F., dos Santos, R.F., Santos, I., Silva, J.P.N., Arraiano, C.M., and Andrade, J.M. (2020). RNA-binding proteins driving the regulatory activity of small non-coding RNAs in bacteria. Front. Mol. Biosci. 7.

Rappsilber, J., Mann, M., and Ishihama, Y. (2007). Protocol for micro-purification, enrichment, pre-fractionation and storage of peptides for proteomics using StageTips. Nat. Protoc. 2: 1896–1906.

Rast, A., Rengstl, B., Heinz, S., Klingl, A., and Nickelsen, J. (2016). The role of Slr0151, a tetratricopeptide repeat protein from Synechocystis sp. PCC 6803, during photosystem II assembly and repair. Front. Plant Sci. 7: 605.

Reimann, V., Alkhnbashi, O.S., Saunders, S.J., Scholz, I., Hein, S., Backofen, R., and Hess, W.R. (2017). Structural constraints and enzymatic promiscuity in the Cas6-dependent generation of crRNAs. Nucleic Acids Res. 45: 915–925.

Rengstl, B., Knoppová, J., Komenda, J., and Nickelsen, J. (2013). Characterization of a Synechocystis double mutant lacking the photosystem II assembly factors YCF48 and Sll0933. Planta 237: 471–480.

Rosana, A.R.R., Denise S. Whitford, Migur, A., Steglich, C., Kujat-Choy, S.L., Hess Wolfgang R., and Owttrim George W. (2020). RNA helicase-regulated processing of the Synechocystis rimO-crhR operon results in differential cistron expression and accumulation of two sRNAs. J Biol Chem 295: 6372–6386.

Sato, H., Fujimori, T., and Sonoike, K. (2008). sll1961 is a novel regulator of phycobilisome degradation during nitrogen starvation in the cyanobacterium Synechocystis sp. PCC 6803. FEBS Lett. 582: 1093–1096.

Schneider, G.J. and Hasekorn, R. (1988). RNA polymerase subunit homology among cyanobacteria, other eubacteria and archaebacteria. J. Bacteriol. 170: 4136–4140.

Scholz, I., Lange, S.J., Hein, S., Hess, W.R., and Backofen, R. (2013). CRISPR-Cas systems in the cyanobacterium Synechocystis sp. PCC6803 exhibit distinct processing pathways involving at least two Cas6 and a Cmr2 protein. PloS One 8: e56470.

Scholz, I., Lott, S.C., Behler, J., Gärtner, K., Hagemann, M., and Hess, W.R. (2019). Divergent methylation of CRISPR repeats and cas genes in a subtype I-D CRISPR-Cas-system. BMC Microbiol. 19: 147.1-147.11.

Schuergers, N., Ruppert, U., Watanabe, S., Nürnberg, D.J., Lochnit, G., Dienst, D., Mullineaux, C.W., and Wilde, A. (2014). Binding of the RNA chaperone Hfq to the type IV pilus base is crucial for its function in Synechocystis sp. PCC 6803. Mol. Microbiol. 92: 840–852.

Schuster, C.F. and Bertram, R. (2013). Toxin-antitoxin systems are ubiquitous and versatile modulators of prokaryotic cell fate. FEMS Microbiol. Lett. 340: 73–85.

Schwarz, D., Schubert, H., Georg, J., Hess, W.R., and Hagemann, M. (2013). The gene sml0013 of Synechocystis sp. strain PCC 6803 encodes for a novel subunit of the NAD(P)H oxidoreductase or complex I that is ubiquitously distributed among Cyanobacteria. Plant Physiol. 163: 1191–1202.

Shah, S.A., Alkhnbashi, O.S., Behler, J., Han, W., She, Q., Hess, W.R., Garrett, R.A., and Backofen, R. (2019). Comprehensive search for accessory proteins encoded with archaeal and bacterial type III CRISPR-cas gene cassettes reveals 39 new cas gene families. RNA Biol. 16: 530–542.

Shmakov, S.A., Makarova, K.S., Wolf, Y.I., Severinov, K.V., and Koonin, E.V. (2018). Systematic prediction of genes functionally linked to CRISPR-Cas systems by gene neighborhood analysis. Proc. Natl. Acad. Sci. 115: E5307–E5316.

Smirnov, A., Förstner, K.U., Holmqvist, E., Otto, A., Günster, R., Becher, D., Reinhardt, R., and Vogel, J. (2016). Grad-seq guides the discovery of ProQ as a major small RNA-binding protein. Proc. Natl. Acad. Sci. 113: 11591–11596.

Spät, P., Macek, B., and Forchhammer, K. (2015). Phosphoproteome of the cyanobacterium Synechocystis sp. PCC 6803 and its dynamics during nitrogen starvation. Front. Microbiol. 6.

Spence, E., Bailey, S., Nenninger, A., Møller, S.G., and Robinson, C. (2004). A Homolog of Albino3/OxaI Is Essential for Thylakoid Biogenesis in the Cyanobacterium Synechocystis sp. PCC6803. J. Biol. Chem. 279: 55792–55800.

Srikumar, A., Krishna, P.S., Sivaramakrishna, D., Kopfmann, S., Hess, W.R., Swamy, M.J., Lin-Chao, S., and Prakash, J.S.S. (2017). The Ssl2245-Sll1130 Toxin-Antitoxin system mediates heat-induced programmed cell death in Synechocystis sp. PCC6803. J. Biol. Chem. 292: 4222–4234.

Summerfield, T.C., Winter, R.T., and Eaton-Rye, J.J. (2005). Investigation of a requirement for the PsbP-like protein in Synechocystis sp. PCC 6803. Photosynth. Res. 84: 263–268.

Sveshnikov, D., Funk, C., and Schröder, W.P. (2007). The PsbP-like protein (sll1418) of Synechocystis sp. PCC 6803 stabilises the donor side of Photosystem II. Photosynth. Res. 93: 101–109.

Trautmann, D., Voss, B., Wilde, A., Al-Babili, S., and Hess, W.R. (2012). Microevolution in cyanobacteria: re-sequencing a motile substrain of Synechocystis sp.P CC 6803. DNA Res. 19: 435–448.

Turmel, M., Gagnon, M.-C., O’Kelly, C.J., Otis, C., and Lemieux, C. (2009). The chloroplast genomes of the green algae Pyramimonas, Monomastix, and Pycnococcus shed new light on the evolutionary history of prasinophytes and the origin of the secondary chloroplasts of euglenids. Mol. Biol. Evol. 26: 631–648.

Uchiyama, I., Mihara, M., Nishide, H., Chiba, H., and Kato, M. (2019). MBGD update 2018: microbial genome database based on hierarchical orthology relations covering closely related and distantly related comparisons. Nucleic Acids Res. 47: D382–D389.

Vijay, D., Akhtar, M.K., and Hess, W.R. (2019). Genetic and metabolic advances in the engineering of cyanobacteria. Curr. Opin. Biotechnol. 59: 150–156.

Vizcaíno, J.A. et al. (2013). The PRoteomics IDEntifications (PRIDE) database and associated tools: status in 2013. Nucleic Acids Res. 41: D1063–1069.

Wagner, E.G.H. and Romby, P. (2015). Small RNAs in bacteria and archaea: who they are, what they do, and how they do it. Adv. Genet. 90: 133–208.

Wassarman, K.M. and Storz, G. (2000). 6S RNA regulates Escherichia coli RNA polymerase activity. Cell 101: 613–623.

Weisz, D.A., Johnson, V.M., Niedzwiedzki, D.M., Shinn, M.K., Liu, H., Klitzke, C.F., Gross, M.L., Blankenship, R.E., Lohman, T.M., and Pakrasi, H.B. (2019). A novel chlorophyll protein complex in the repair cycle of photosystem II. Proc. Natl. Acad. Sci. U. S. A. 116: 21907–21913.

Westermann, A.J., Venturini, E., Sellin, M.E., Förstner, K.U., Hardt, W.-D., and Vogel, J. (2019). The major RNA-binding protein ProQ impacts virulence gene expression in Salmonella enterica serovar Typhimurium. mBio 10: e02504–18.

Winkelman, J.T., Blair, K.M., and Kearns, D.B. (2009). RemA (YlzA) and RemB (YaaB) regulate extracellular matrix operon expression and biofilm formation in Bacillus subtilis. J. Bacteriol. 191: 3981–3991.

Winkelman, J.T., Bree, A.C., Bate, A.R., Eichenberger, P., Gourse, R.L., and Kearns, D.B. (2013). RemA is a DNA-binding protein that activates biofilm matrix gene expression in Bacillus subtilis: RemA activates matrix genes directly. Mol. Microbiol. 88: 984–997.

Wower, I.K., Zwieb, C.W., Guven, S.A., and Wower, J. (2000). Binding and cross-linking of tmRNA to ribosomal protein S1, on and off the Escherichia coli ribosome. EMBO J. 19: 6612–6621.

Xu, C., Wang, B., Yang, L., Hu, L.Z., Yi, L., Wang, Y., Chen, S., Emili, A., and Wan, C. (2020). Global landscape of native protein complexes in Synechocystis sp. PCC 6803. bioRxiv: 2020.03.07.980128.

Yamaguchi, K. and Subramanian, A.R. (2003). Proteomic identification of all plastid-specific ribosomal proteins in higher plant chloroplast 30S ribosomal subunit. Eur. J. Biochem. 270: 190–205.

Zhang, A., Altuvia, S., Tiwari, A., Argaman, L., Hengge-Aronis, R., and Storz, G. (1998). The OxyS regulatory RNA represses rpoS translation and binds the Hfq (HF-I) protein. EMBO J. 17: 6061–6068.

Zhang, B. and Horvath, S. (2005). A general framework for weighted gene co-expression network analysis. Stat. Appl. Genet. Mol. Biol. 4.

Zhang, J., Gao, F., Zhao, J., Ogawa, T., Wang, Q., and Ma, W. (2014a). NdhP is an exclusive subunit of large complex of NADPH dehydrogenase essential to stabilize the complex in Synechocystis sp. strain PCC 6803. J. Biol. Chem. 289: 18770–18781.

Zhang, J.-Y., Deng, X.-M., Li, F.-P., Wang, L., Huang, Q.-Y., Zhang, C.-C., and Chen, W.-L. (2014b). RNase E forms a complex with polynucleotide phosphorylase in cyanobacteria via a cyanobacterial-specific nonapeptide in the noncatalytic region. RNA N. Y. N 20: 568–579.

Zheng, J.J., Perez, A.J., Tsui, H.-C.T., Massidda, O., and Winkler, M.E. (2017). Absence of the KhpA and KhpB (JAG/EloR) RNA-binding proteins suppresses the requirement for PBP2b by overproduction of FtsA in Streptococcus pneumoniae D39. Mol. Microbiol. 106: 793–814.

## Supplemental references

Oberto, J. (2013). SyntTax: a web server linking synteny to prokaryotic taxonomy. BMC Bioinformatics 14: 4.

Yang, H., Liu, J., Wen, X., and Lu, C. (2015). Molecular mechanism of photosystem I assembly in oxygenic organisms. Biochim. Biophys. Acta BBA - Bioenerg. 1847: 838–848.

